# T cell receptor and IL-2 signaling strength control memory CD8^+^ T cell functional fitness via chromatin remodeling

**DOI:** 10.1101/2021.10.19.464978

**Authors:** Shu Shien Chin, Erik Guillen, Laurent Chorro, Sooraj Achar, Karina Ng, Susanne Oberle, Francesca Alfei, Dietmar Zehn, Grégoire Altan-Bonnet, Fabien Delahaye, Grégoire Lauvau

## Abstract

Cognate antigen signal controls CD8^+^ T cell priming, expansion size and effector versus memory cell fates, but it is not known if and how it modulates the functional features of memory CD8^+^ T cells. Here we show that the strength of T cell receptor (TCR) signaling determines the requirement for interleukin-2 (IL-2) signals to form a pool of memory CD8^+^ T cells that competitively re-expand upon secondary antigen encounter. Combining strong TCR and intact IL-2 signaling synergistically induces genome-wide chromatin accessibility in regions targeting a wide breadth of biological processes, consistent with their greater functional fitness. Chromatin accessibility in promoters of genes encoding for stem cell, cell cycle and calcium-related proteins correlated with faster intracellular calcium accumulation, initiation of cell cycle and more robust expansion. High-dimensional flow-cytometry analysis also highlights higher subset diversity and phenotypes. These results formally establish that epitope selection in vaccine design strongly impacts memory CD8^+^ T cell epigenetic programming and functions.

**One Sentence Summary:** The strength of antigenic and interleukin 2 signals received by CD8^+^ T cells during vaccination epigenetically programs their ability to form functional memory.

## Introduction

Most currently approved vaccines in humans rely on strong antibody (Ab)-mediated immunity and only few trigger effective CD8^+^ T cell immunological memory ^1^. CD8^+^ T cells can uniquely detect intracellular antigens and are equipped to directly kill microbial pathogen-infected and tumor cells. CD8^+^ T cells are also less prone to be impacted by point mutations that alter conformation sensitive Ab epitopes. Therefore, eliciting a pool of long-lived functional memory CD8^+^ T cells through rational design remains a necessity ^2, 3^.

Single-cell tracing studies have established that one CD8^+^ T cell has the potential to give rise to multiple progenies, supporting the idea that as T cells expand from an original clone, they differentially integrate priming signals, leading to distinct phenotypic fates ^4, 5^. While the initiation of clonal expansion most likely results from a digital on-off response, the functional fates of T cells are rather the consequence of an analog response to the sum of the different priming signals ^6–9^. The current dogma states that the greater the T cell receptor (TCR) signals are, the stronger the primary expansion of CD8^+^ T cells is, and the more effector cells are produced at the expense of memory cells ^10, 11^. While cytokines, e.g., IL-2, IL-12, type I IFN, strongly skew CD8^+^ T cell-differentiation towards robust effector cells by directly enhancing expression of transcriptional regulators such as T-bet and Blimp-1 in the T cells ^12–17^, cytokines can also directly potentiate TCR signaling ^18^. When TCR signaling augments, the overall number of antigen-specific effector and memory CD8^+^ T cells that are generated also increases, but memory cells primed with different strengths of TCR signals do not differ in their ability to clonally re-expand *in vivo* ^19^. Memory CD8^+^ T cells primed with weak TCR signals, however, fail to express effector functions in response to weak (but not strong) epitopes, suggesting that weak TCR signals during priming may induce functionally distinct memory cells ^20^. Recent work investigating T cell epigenetic programming further established that the acquisition of functional effector or memory features by CD8^+^ T cells is achieved through specific histone modification-driven chromatin remodeling that occurs at early/effector stages of T cell differentiation ^21–26^. In these studies, a significant overlap in the epigenetic landscapes between effector and memory CD8^+^ T cells has generally been reported. Although the memory CD8^+^ T cell epigenetic landscape is largely distinct from that of naïve T cells, memory cells also reacquire part of the naïve T cell-associated landscape via active demethylation of genes associated with naïve T cells status ^23^.

Despite many studies investigating the steps orchestrating CD8^+^ T cell priming and differentiation into memory cells, there is still an acute gap in knowledge as to whether the strength of TCR and cytokine signals modulate the functional features and the epigenetic programming of memory CD8^+^ T cells. In this work, we tested the hypothesis that the strength of TCR signals regulates naïve CD8^+^ T cell dependency on IL-2 signals and their differentiation into functional memory cells. We next determined whether altering the strength of these signals individually or simultaneously modulates memory CD8^+^ T cell transcriptomic and epigenetic programming.

## Results

### TCR signaling strength modulates early CD8^+^ T cell activation and transcriptional program

To investigate whether TCR signaling strength alters early naïve CD8^+^ T cell activation, we used two altered peptide ligands (APLs) of the ovalbumin-derived K^b^-presented SIINFEKL epitope (Ova257-264 or “N4”) that both exhibit low potency for stimulating OT-I transgenic T cells compared to the natural N4 epitope. One variant (“A8”) decreases peptide/MHC stability after substituting the C-terminus leucine anchor residue with alanine while the other (“T4”) alters OT-I TCR/peptide interactions upon replacement of the asparagine by a threonine (Figure S1A). T4 and N4 both stabilize K^b^ similarly whereas A8 is a much weaker K^b^ stabilizer (by a factor of ∼3- 4, Figure S1B ^19, 27^). Quantitative multi-parameter kinetic monitoring of OT-I cells stimulated *in vitro* with N4, A8 and T4 peptides revealed that the proportion of OT-I cells undergoing at least one division (Fraction diluted) and the extent of their expansion (Proliferation index) was substantially lower for OT-I cells that received weak TCR signals compared to those that were stimulated with strong TCR signals (Figure 1A and Figure S1C). This was observed across a wide range of peptide concentrations (from 10^-6^ to 10^-9^ M) and as a result of low epitope/MHC stability (A8) or weak OT-I TCR interaction with its cognate epitope (T4). Moreover, such weakly-stimulated (A8, T4) OT-I cells secreted little to no cytokines (IL-2, IFNγ, TNFα) and only expressed low levels of the Th1 master transcription factor T-bet. Overall, these *in vitro* data show a strong impact of the strength of TCR signals on CD8^+^ T cell proliferation/expansion, cytokine secretion and effector differentiation. To achieve a more global understanding of how naïve CD8^+^ T cells integrate the distinct levels of TCR signals *in vivo*, we next FACS-sorted activated/divided (CFSE^low^) OT-I cells 3 days post-infection with *Listeria monocytogenes (Lm)* expressing either A8 or T4 APLs, or the N4 epitope and conducted whole genome expression arrays (Figure 1B and Figure S1D). Principal component analysis (PCA) shows that, as expected, OT-I cells primed with weak (A8, T4) or strong (N4) TCR signals segregated away from naïve counterparts. A8- and T4-primed OT-I cells clustered together but apart from N4-counterpart, in line with our *in vitro* quantitative analysis where A8- and T4-primed OT-I cells underwent similar phenotypic and functional changes. This result was also reflected in the large overlap of genes differentially expressed in A8- or T4- compared to N4-primed OT-I cells (∼800 genes).

**Figure 1.**
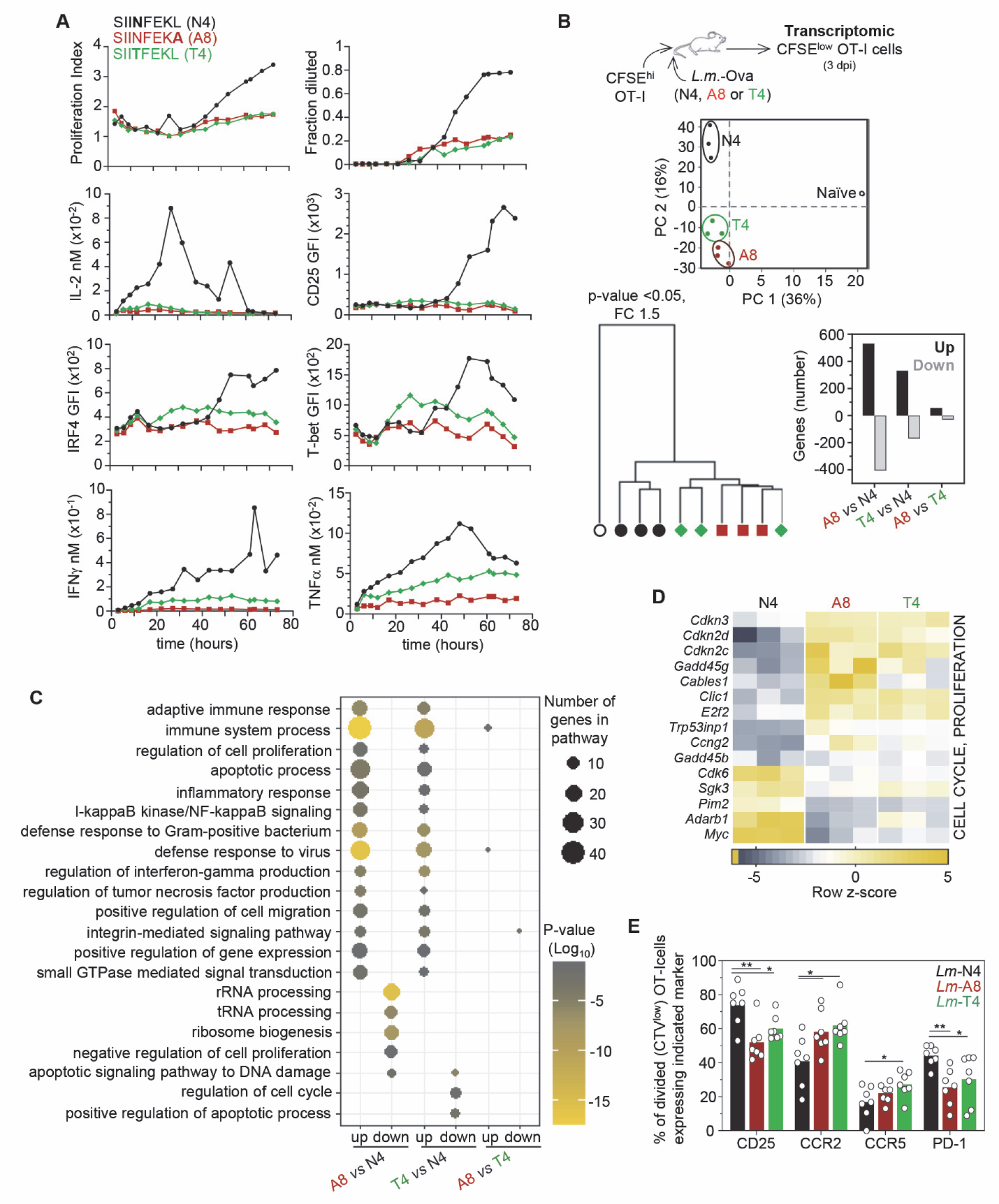
Impact of TCR signaling strength on early CD8^+^ T cell activation. (**A**) Naïve OT-I cells from spleens were stained with CFSE and stimulated *in vitro* with 10^-9^ or 10^-10^ M of ovalbumin (Ova) peptide SIINFEKL (N4) or its altered peptide ligands (T4, A8) for quantitative multi-parameter kinetic monitoring. OT-I cells were stained for cell surface CD25 and intracellular IRF4 and T-bet, and secreted IL-2, IFNγ and TNFα were quantified as described. Each graph summarizes the extend of OT-I cell division (Proliferation index and Fraction diluted at 10^-10^ M of peptides), OT-I cell expression levels (GFI) of stained markers or secreted cytokine levels (at 10^-9^ M of peptides). (**B**) Naïve spleen-derived CFSE-labelled OT-I cells were adoptively transferred to recipient mice, subsequently infected with *Lm-*Ova N4, A8 or T4. 3 days later, CFSE^low^ OT-I cells were flow-sorted from spleens of 3 independent replicates of mice. Total RNA was extracted, reverse transcribed to cDNA before running an Affymetrix mouse expression arrays (Pico 1.0). Principal component analysis (PCA) and hierarchical clustering of expressed genes in analyzed groups with each symbol featuring one mouse. Bar graph shows significantly up- and down-regulated genes, with fold change +/-1.5 and *p*-value≤ 0.05. **(C)** Network analysis of biological-process gene-ontology (GO) term enrichment among significantly up- or down-regulated genes in the indicated priming comparisons. Differentially regulated genes were analyzed for over-represented GO terms on DAVID website and visualized in a scatterplot graph. Node color is proportional to the FDR-adjusted *p*-value of the enrichment. Node size is proportional to the number of genes in each GO term. **(D)** Heat map of cell- cycle/proliferation-related genes for which expression is significantly different between N4, A8 and T4 primed OT-I cells. Row *z*-score fold change is shown in heat map. **(E)** Proportion of CTV^low^ (divided) OT-I cells expressing indicated markers 3 days post immunization with *Lm-* Ova N4, A8 or T4.

GO pathway analysis of upregulated genes in OT-I cells primed with *Lm* expressing A8 or T4 compared to N4 revealed an overlap in pathways related to the immune response (adaptive, inflammation, interferon and tumor necrosis factor, defense to virus/bacteria), cell cycle/proliferation and death, adhesion and migration (Figure 1C, Table S1). Differences were noted in the pathways related to the differentially downregulated genes but these pathways were mostly involved in non-immune functions. Essentially no pathways were found when comparing A8- and T4-primed OT-I cells, consistent with the very small number of differentially expressed genes between these two priming conditions. Among the differentially induced genes between A8 or T4- versus N4-primed OT-I cells, many encoded cell-cycle-cyclin-dependent kinases (*Cdkn3, Cdkn2d, Cdkn2c, Cables1, Cdk6*) and transcription factors (TFs) (*Myc, Irf4, Mxd1, E2f2*), interferon-related proteins (*Ifit1, Ifit2, Isg15, Oas1a, Oas3, Ddx58, Nod1*) and TFs (*Irf7, Stat2*), migration and chemotaxis (*Itgax, Sell, Ccr2, Ccr5, S1p1r, klf2*) and effector functions (*Gzma, Gzmk, Klrc1, Klrd1, Klrc2, Klrc3*) (Figures 1D and S1E). Several of these genes were upregulated in A8- and T4- compared to N4-primed OT-I cells, indicating that naïve CD8^+^ T cells receiving weak TCR signals overall incorporate greater signals related to the highlighted pathways. For instance, expression of several genes coding for proteins involved in cell cycle inhibition (*Cdkn3, Cdkn2d, Cdkn2c, Cables1, E2f2, Trp53inp1)* was increased in OT-I cells primed with the weak A8 and T4 APLs while others implicated in robust cell cycle progression were diminished (*Myc, Cdk6, Irf4*) compared to OT-I cells primed with the strong N4 epitope.

Greater upregulation of the high affinity IL-2 receptor alpha chain CD25 on divided (CTV^low^) OT-I cells was also confirmed by FACS 3 days post *Lm*-N4 immunization (Figure 1E), consistent with prolonged proliferative program compared to A8- and T4-primed counterparts. Thus, T cell proliferation, a major readout outcome in OT-I cells primed with the N4, T4 or A8 epitope *in vitro* and *in vivo* seemed consistent. We also tested whether other differentially expressed genes in activated dividing OT-I cells translated into distinct protein expression levels 3 days post immunization (Figures 1E and S1F). We confirmed findings for CCR2, CCR5 and PD-1 which also reflected different activation states of primed T cells. However, expression of CD127, KLRD1 or CD11c was not significantly distinct possibly reflecting differences in the kinetics of gene versus protein expression. Collectively, modulating the amount of TCR signals incorporated by a naïve CD8^+^ T cell clone during priming *in vitro* and *in vivo* drives a clearly distinct but consistent functional and transcriptional activation program.

### TCR signaling strength determines the requirement for IL-2-signals to form functional memory

Since OT-I cells that received weak compared to strong TCR stimulation exhibited less sustained *in vitro* proliferation, and upregulated genes involved in shutting off cell cycle *in vivo* (Figure 1), we further explored how altering IL-2 signaling, the key T cell proliferative signal, affected T cell clonal expansion and functional memory cell formation in the context of distinct strengths of TCR signals. We hypothesized that in situations of weak TCR stimulation and impaired IL-2 signaling, CD8^+^ T cells may not form functional memory cells. To test this hypothesis, we used *Il2ra^mut/mut^* OT-I cells, that are impaired in IL-2 signaling as a result of a point mutation in an evolutionarily conserved tyrosine (Y129) ^28^. *Il2ra^mut/mut^* OT-I cells were co-transferred with WT counterparts into recipient mice subsequently infected with *Lm* expressing distinct Ova APLs (Q4, T4 or A8) or the N4 epitope (Figure 2A). As expected ^19^, primary expansion of OT-I cells (day 7) was significantly reduced by lowering TCR signaling, and OT-I cells impaired in IL-2 signaling (*Il2ra^mut/mut^* ) proliferated 3 times less than WT counterparts whether infected with any of the *Lm*-Ova APLs (Figure S2A). At memory stage, i.e., >40 days post primary *Lm* infection, we sorted and co-transferred the same number (1,000) of *Il2ra^mut/mut^* and WT OT-I memory cells to naïve mice, that were subsequently challenged with *Lm*-N4 to rigorously assess their ability to competitively expand 6.5 days later (Figure 2A). While N4-primed *Il2ra^mut/mut^* and WT OT-I memory cells underwent comparable proliferation, Q4-, T4- and A8-primed *Il2ra^mut/mut^* OT-I memory cells did not and expanded ∼4-5 times less than WT counterparts (Figure 2B).

**Figure 2.**
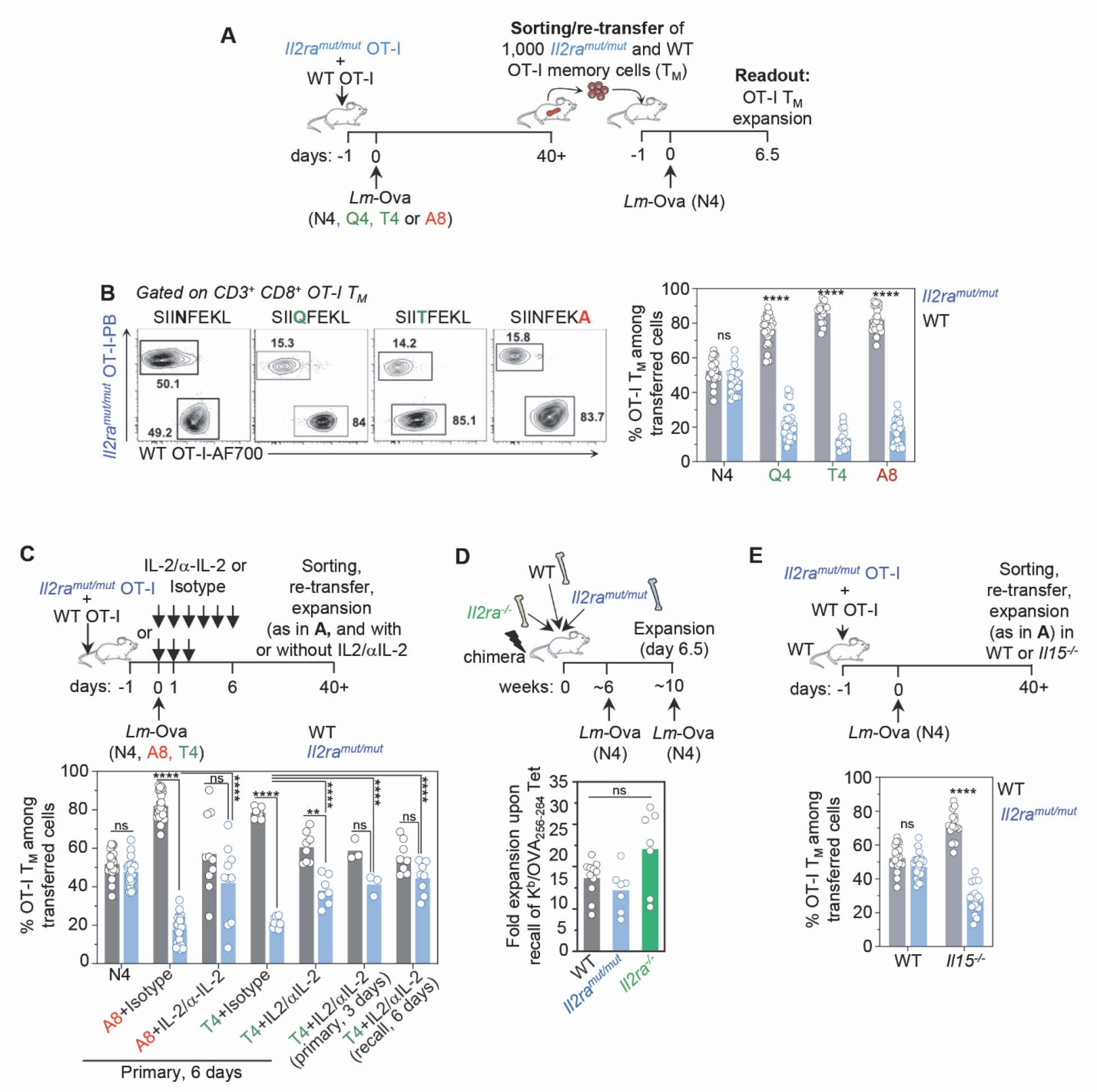
CD8^+^ T cells primed with weak TCR and IL-2 signals failed to form memory cells that competitively expand during recall infection. (**A**) Schematic of experimental design used or referred to in B, C, E. Briefly, 2,000 *Il2ra^mut/mut^* or WT OT-I cells expressing constitutive tomato (Td) and congenically distinct, were adoptively transferred to recipient mice, and infected the next day with *Lm*-Ova N4, Q4, T4 or A8. *Il2ra^mut/mut^* and WT OT-I memory cells (∼day 40 p.i.) were then sorted by FACS and transferred at a 1:1 ratio to new recipient mice subsequently infected with *Lm*-OvaN4 the next day. 6.5 days later, spleens were harvested and stained to quantify *Il2ra^mut/mut^* and WT OT-I memory cell expansion. (**B**) Following (A), representative FACS dot plots of re-expanded *Il2ra^mut/mut^* and WT OT-I memory cells originally primed with the indicated Ova APLs are shown. Bar graph represents the pooled frequencies of re-expanded *Il2ra^mut/mut^* versus WT OT-I memory cells across >6 independent replicate experiments (n=30 mice). (**C**) As in (A), but mice were also injected with IL-2/anti-IL2 mAb complexes during initial or recall infection, every day for 6 or 3 days as depicted on the schematic and in Figure S2B. The bar graph shows the relative average frequency of re-expanded *Il2ra^mut/mut^* and WT OT-I memory cells across 1-4 independent replicate experiments (n=3-30 mice). (**D**) WT/*Il2ra^mut/mut^*/*Il2ra^-/-^* mixed BM chimeras (ratio 1:1:1) were infected i.v. with *Lm*-OvaN4 and challenged 4 weeks later with *Lm*-OvaN4. 6.5 days later, splenocytes were stained with Ova257- 264/K^d^ tetramers (Tet^+^) and appropriate congenic markers. Bar graphs show the relative fold expansion of Tet^+^ CD8^+^ T cells across 2-3 independent replicate experiments (n=11 mice). (**E**) As in (A), but FACS-sorted *Il2ra^mut/mut^* and WT OT-I memory cells were adoptively transferred to either WT or *Il15^-/-^* hosts. The bar graph shows the summary of the relative frequency of re- expanded *Il2ra^mut/mut^* and WT OT-I memory cells across 4-5 replicate experiments (n=17-24 mice). In all panels, each symbol represent 1 mouse and *p*-values are indicated when relevant with \**p* < 0.05; \*\**p*< 0.01; \*\*\**p*< 0.001; \*\*\*\**p*< 0.0001; NS, not significant, using two-tailed unpaired Student’s *t* test.

Consistent with our hypothesis, this result indicated that TCR signaling strength determines the dependency of naïve CD8^+^ T cells on intact IL-2 signaling for their differentiation into memory cells capable of competitive clonal expansion. Providing robust IL-2 signals to *Lm*-A8- and *Lm*- T4-primed *Il2ra^mut/mut^* OT-I cells, using IL-2/anti-IL-2 monoclonal Ab (mAb) treatment for 6 days during primary infection, showed significant rescue of their ability to expand compared to control isotype Ab-treated counterparts during secondary antigen encounter after *Lm-*N4 challenge infection (Figure 2C). IL-2-rescued *Il2ra^mut/mut^* OT-I memory cells primed with the weak Ova APLs (A8, T4) expanded closely to WT OT-I memory cells. When IL-2 signals were given for only 3 instead of 6 days, the extent of the rescue was comparable, suggesting that IL-2 signals were mostly important at early stages post-priming. Interestingly, if IL-2 signals were only provided during the recall response, *Il2ra^mut/mut^* OT-I memory cells primed with weak APL (here T4) expanded like WT counterparts, indicating that providing robust IL-2 signaling during Ag re-encounter also restores their ability to competitively expand (Figure 2C and S2B). To extend results to polyclonal memory CD8^+^ T cells, we next tracked *Il2ra^mut/mut^*, *Il2ra^-/-^* and WT N4-primed polyclonal K^b^/Ova254-261 tetramer^+^ (Tet^+^) memory CD8^+^ T cells in mixed bone marrow (BM) chimera mice reconstituted with an equal ratio of each of the above BM genotype (Figure 2D). We confirmed that antigen-specific memory CD8^+^ T cell of all genotypes proliferated independently of IL-2 upon secondary Ag encounter. However, when N4-primed *Il2ra^mut/mut^* and WT OT-I memory cells were transferred to *Il15^-/-^* mice, they failed to expand equivalently after challenge with either *Lm* or VSV expressing OvaN4, indicating that STAT5 signaling downstream of either IL-15 or IL-2 cytokine stimulation is essential for maximal clonal memory cell expansion (Figure 2E and Figure S2C, D). In summary, these results establish that the strength of TCR signaling regulates CD8^+^ T cell-dependency on IL-2 signals in order to form memory cells that can effectively expand during secondary Ag encounter. The data further suggest that robust IL-2 signals are mostly required early on post CD8^+^ T cell priming, but that IL-2 signals provided during the recall response can also rescue memory CD8^+^ T cell competitive expansion.

### Existence of IL-2-dependent and IL-2-independent natural epitopes to form functional memory

We next investigated whether naturally presented epitopes may also require IL-2 signaling during priming in order to form functional memory cells that competitively expand during a recall infection. We picked five epitopes expressed in *Lm*, all originally derived from three distinct microbial pathogens, the lymphochorionmeningitis virus (LCMV, GP33-41 and GP276-284), the herpes simplex virus 2 (HSV-2, gB498-505) and *Lm* (LLO91-99 and p60217-225)(Figure 3A). We first generated mixed bone marrow (BM) chimeras in lethally irradiated WT recipient mice reconstituted with BM from either *Il2ra^mut/mut^*, *Il2ra^-/-^* and WT mice or from *Il2ra^mut/mut^* and WT mice. We used BM donor and recipient mice that express i) distinct CD45 congenic marker combinations on the C57BL/6 genetic background (H-2^b^) and ii) the K^d^ molecule (B6-K^d^). Fully reconstituted chimeras were next primary infected and further challenged with the same microbial pathogen (autologous) 4-8 weeks later (Figure 3B). Fold expansion of polyclonal Tet^+^ memory CD8^+^ T cells of each genotype and specific for each listed epitope was then quantified by comparing the average splenic frequencies of Tet^+^ memory cells prior to secondary challenge to those 6.5 days post challenge (Figure 3C, D). While *Il2ra^-/-^* or *Il2ra^mut/mut^* GP33-41/D^b^ and GP276-284/D^b^-specific memory CD8^+^ T cells failed to expand as competitively as WT counterparts, and required robust IL-2 signals during priming for competitive secondary expansion, those recognizing gB498-505/K^b^, LLO91-99/K^d^ and p60217-225/K^d^ did not (Figure 3C).

**Figure 3.**
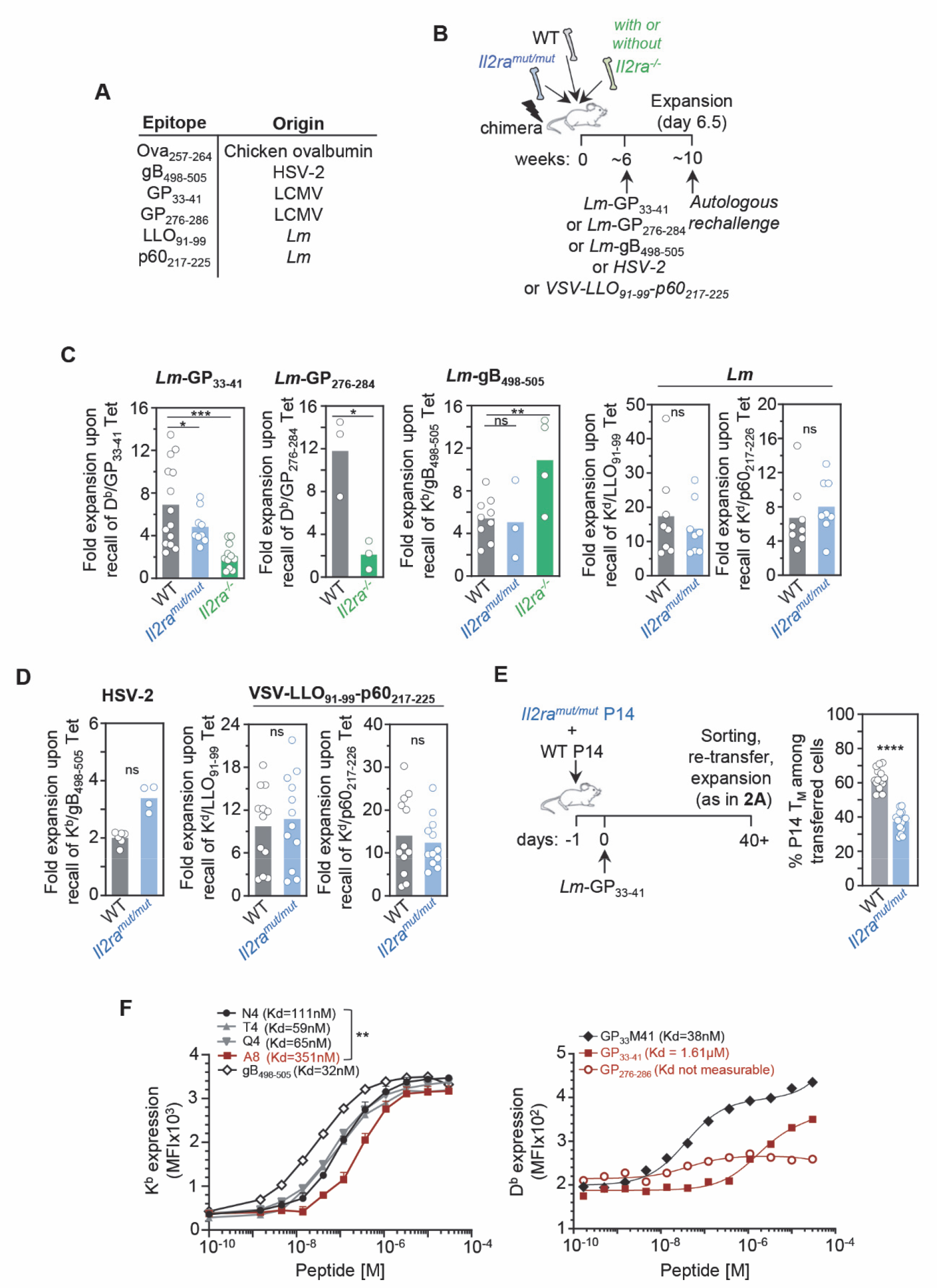
Several natural epitopes require IL-2 signaling to form functional memory CD8^+^ T cells. **A** Table listing the naturally presented epitopes used in our studies and their microbial pathogen origin. **(B)** Schematic of experimental design for experiments in C, D. (**C**) WT/*Il2ra^mut/mut^* or WT/*Il2ra^mut/mut^*/*Il2ra^-/-^* mixed BM chimeras (ratio 1:1:1) were infected i.v. with indicated pathogens 6-8 weeks post-reconstitution, and challenged with the same autologous pathogen 4-5 weeks after before staining for FACS analysis 6.5 days later using indicated MHC class I tetramers (Tet^+^) and appropriate congenic markers. Bar graphs show the relative fold expansion of Tet^+^ CD8^+^ T cells across 2 replicate experiments (n=8-15 mice). **(D)** WT/*Il2ra^mut/mut^* mixed BM chimeras were set up as in (B), infected and then challenged 4 weeks later with indicated pathogens before staining spleen cells for FACS analysis 4 or 6.5 days later. Bar graphs show the relative fold expansion of Tet^+^ CD8^+^ T cells across 2 replicate experiments (n=5-12 mice). **(E)** *Il2ra^mut/mut^* and WT P14 cells were adoptively transferred to naive hosts, subsequently infected the next day with *Lm*-GP33-41. *Il2ra^mut/mut^* and WT P14 memory cells were sorted 4 weeks later, transferred (ratio 1:1) to naive hosts, and challenged the next day with *Lm*- GP33-41. Spleens cells were stained for FACS analysis 6.5 days later. Bar graph shows the relative frequency of *Il2ra^mut/mut^* and WT P14 memory cells across 3 replicate experiments (n=15 mice). (**F**) H2-K^b^ and H2-D^b^ RMA-S stabilization assay by indicated peptides. Kd values represent the peptide concentration required to achieve half of maximum cell-surface K^b^ or D^b^ expression across 3 independent replicate experiments. Each symbol (C-E) represents one mouse and *p*- values are indicated when relevant with \**p* < 0.05; \*\**p*< 0.01; \*\*\**p*< 0.001; \*\*\*\**p*< 0.0001; NS, not significant, using two-tailed unpaired Student’s *t* test.

Next, using HSV-2 or VSV-expressing the *Lm*-derived LLO91-99 and p60217-225 epitopes to infect and challenge *Il2ra^mut/mut^*/WT mixed BM chimeras, we revealed that these epitopes also did not depend on IL-2 to form functional memory cells, similarly to when they were expressed in *Lm* (Figure 3D). Using the clonal population of GP33-41/D^b^-specific *Il2ra^mut/mut^* and WT P14 TCR transgenic memory CD8^+^ T cells primed upon *Lm*-GP33-41 infection, we further confirmed that P14 memory CD8^+^ T cells also failed to competitively expand when IL-2 signals were impaired (Figure 3E). Notably, during primary infection of the chimera mice, some epitopes (Ova257-264, gB498-505) but not others (GP33-41) required IL-2 signals for optimal expansion, whether expressed in *Lm* or by other microbial pathogens (VSV, HSV-2)(Figure S3B-G). However, the dependency on IL-2 signals for maximal primary expansion did not predict that of the recall response. Since these various microbial pathogen infections are likely to induce a very distinct inflammatory environment in infected hosts, this result further suggested that IL-2 dependency for formation of functional memory is more likely to be epitope-driven rather than inflammation-driven.

Consistent with such interpretation, the GP33-41 and GP276-286 epitopes, failed to form stable complexes with D^b^ in *in vitro* RMA-S D^b^ stabilization assay, compared to the GP33M41 peptide, a stabilized version of the GP33-41 epitope (Figure 3F). Likewise, the gB498-505 epitope, like N4 but in contrast to A8, was not dependent on IL-2 signaling, and could efficiently stabilize K^b^. Thus, epitopes that cannot form stable complexes with their respective MHC may only trigger weak TCR signaling (like T4 and Q4 APLs), and be dependent on IL-2 signals to induce functional memory cells. Taken together, these results suggest that the amount of TCR signal an epitope gives to naïve polyclonal CD8^+^ T cells, but not a specific set of T cell clones or inflammatory factors elicited by distinct microbial pathogen infections, will most likely determine the functional fates of memory CD8^+^ T cells and their dependency on cytokine signals. These data also indicate that natural epitopes can trigger distinct memory CD8^+^ T cell programs that either did or did not require intact IL-2 signals to form a pool of fully functional memory cells.

### The duration of cognate T cell antigen presentation may not dictate the formation of functional memory

To explore if the duration of epitope presentation *in vivo* correlates with the induction of functional memory CD8^+^ T cells, we adoptively transferred CTV-labelled OT-I, P14, L9.6 (p60217-225/K^d^-specific) and gBT-I (gB498-505/K^b^-specific) CD8^+^ T cells in mice infected with *Lm* expressing N4, T4, A8 or either of the other epitopes 1, 3 or 5 days before (Figure 4A). While both Ova (N4)- and gB498-505-derived epitopes could respectively prime naïve OT-I and gBT-I T cells and induce their robust proliferation for at least five days *in vivo*, *Lm*-A8, LCMV-GP33-41- and *Lm* p60217-225-derived epitopes only induced OT-I, P14 and L9.6 T cell proliferation for ∼2 days, respectively. Since CD8^+^ T cells specific for Ova(N4)-, gB- and p60-derived epitopes did not require IL-2 signals to form functional memory but those specific for GP33-41 or induced by A8 did (Figure 3C, D), it further suggested that the length of epitope presentation *in vivo* and the formation of functional memory cells were uncoupled. Consistent with this finding, OT-I memory cells were primed by the weak T4 APL for at least 3 days, although robust IL-2 signals were required to form functional memory cells.

**Figure 4.**
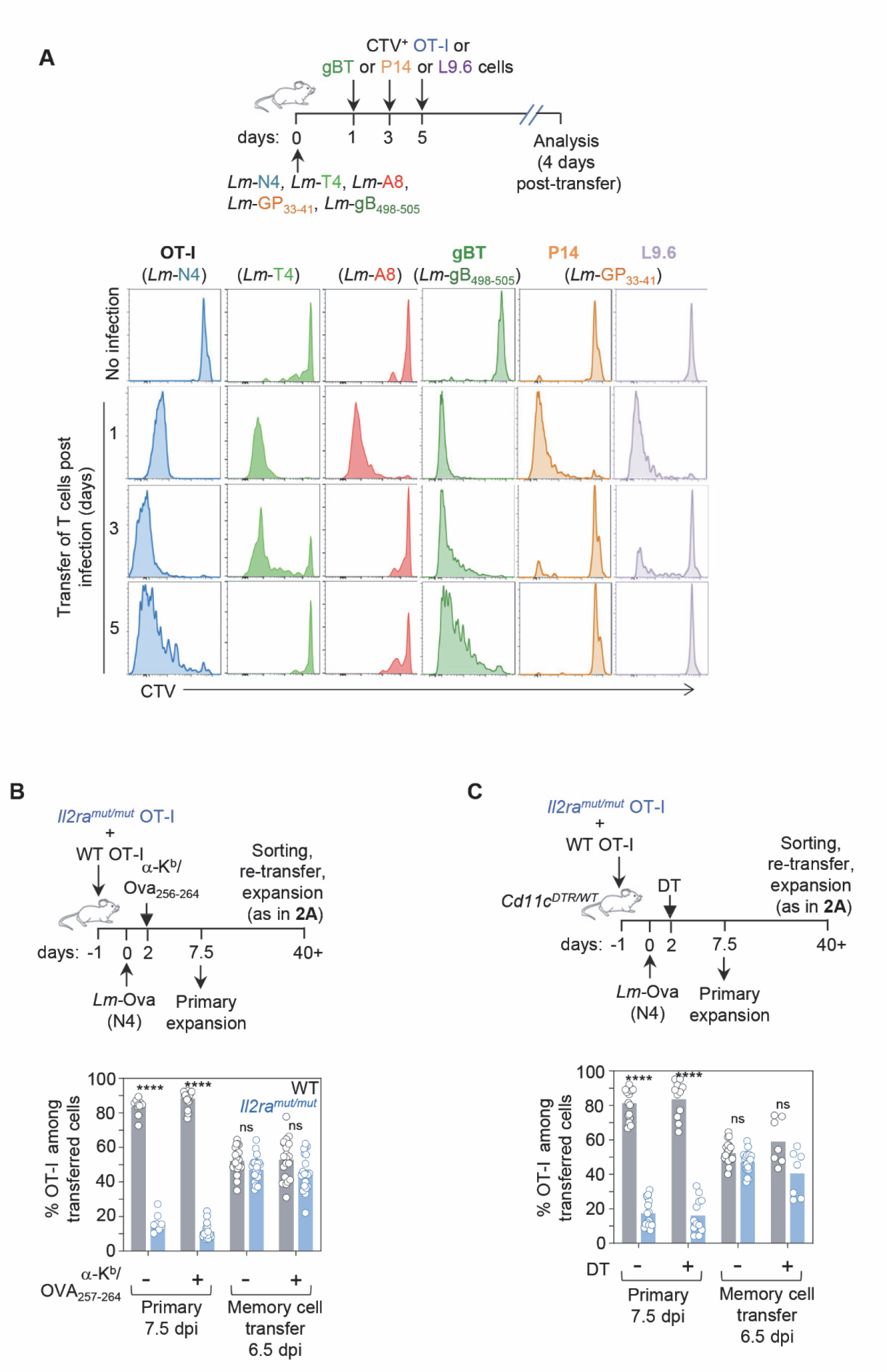
The duration of cognate antigen presentation and the formation of functional memory CD8^+^ T cells are uncoupled. **A** WT mice were infected with indicated *Lm* strains. At specified times, OT-I, gBT, P14 or L9.6 cells were labelled *ex vivo* with CTV and adoptively transferred to previously infected hosts. Spleen cells were stained for FACS analysis 4 days later. Representative FACS histograms show the dilution of CTV staining in transferred T cells. (**B**) *Il2ra^mut/mut^* and WT OT-I cells adoptively transferred to naïve WT mice were subsequently infected the day after with *Lm*-Ova N4. Antigen presentation was disrupted by injecting anti- MHC-I K^b^/Ova256-264 mAb 48 hrs post-infection and spleen cells were stained for FACS analysis either 7.5 days later or 6.5 days after re-transfer of sorted memory cells (as in Figure 2A). Bar graph shows the relative frequency of *Il2ra^mut/mut^* versus WT OT-I cell expansion in 2-5 replicate experiments (n=10-24 mice). (**C**) Same design as in (B), but with *Il2ra^mut/mut^* and WT OT-I cells adoptively transferred to CD11c^DTR/WT^ recipient mice further injected with diphtheria toxin (DT) 48 hours post-infection to deplete CD11c^+^ cells. The bar graph shows the relative frequency of *Il2ra^mut/mut^* versus WT OT-I cell expansion across 2-4 replicate experiments (n=7-16 mice) and each symbol represents one mouse.

To further assess this hypothesis, we next tested if disrupting epitope presentation 2 days after priming indeed prevented the induction of functional memory cells. Mice were co-transferred with *Il2ra^mut/mut^* and WT OT-I cells and either treated with an Ova257-264/K^b^ blocking mAb (WT mice) or selectively depleted of dendritic cells (DCs) upon diphtheria toxin-injection (*Cd11c^DTR/WT^* mice) two days post *Lm*-Ova immunization (Figure 4B, C). Primary OT-I T cell expansion in the blood was decreased by ∼40-50%, confirming efficacy of both treatments (Figure S4A, B). CD11c^+^ cells were depleted in DT-treated *Cd11c^DTR/WT^* mice, and when DT was given prior immunization, OT-I T cells failed to expand (Figure S4C ^29^). Six weeks post immunization, OT-I memory cells (1,000) from treated or control groups were sorted and co-transferred to recipient mice subsequently challenged with *Lm*-N4. OT-I memory cell expansion quantified 6.5 days later was similar whether epitope presentation, by either of the outlined methods, was disrupted or not. In these experimental settings, cognate Ag presentation by non-DCs, or as a result of insufficient mAb blockade, may however, still occur. Nevertheless, these data and the fact that both GP33-41- and p60217-225-derived epitopes were only presented for a short duration of time but exhibited distinct requirements on IL-2 signals to form functional memory, were consistent with a model in which the strength of initial TCR signaling, but not the duration of cognate Ag presentation, is most likely to determine whether naïve CD8^+^ T cells will depend on IL-2 signals to form fully functional memory cells.

### Varying TCR and IL-2 signaling strength strongly modify memory CD8^+^ T cell chromatin accessibility but not transcriptomic profiles

To gain a deeper understanding of the general mechanism by which different strengths of TCR and IL-2 signals shape memory CD8^+^ T cell programming, we next sought to determine and compare the transcriptional and chromatin accessibility profiles of *Il2ra^mut/mut^* and WT OT-I memory CD8^+^ T cells induced upon weak (T4) or strong (N4) *Lm*-expressed epitope priming.

Four weeks post-infection, OT-I memory cells from each of the four experimental groups were flow-sorted and whole genome analysis of both gene expression (by RNA-seq) and open chromatin regions (OCR) (by ATAC-seq) were conducted (Figure 5, Figure S5 and Table S2). Only 28 to 69 genes out of a total of 47, 643 genes (i.e., 0.05-0.15%) were differentially expressed at least 1.5 fold (adjusted *p*-value ≤0.05) when comparing gene expression between *Il2ra^mut/mut^* and WT memory CD8^+^ T cells primed with weak or strong epitope (Figure S5A, B and Table S2). Very few differentially expressed genes were found across the various comparisons when only TCR (e.g., *cdc42ep2*) or IL-2 (e.g., *Eno1b*) varied but a larger number were found when both signals were concomitantly modulated (e.g., *Il27, Tnfsf4, Ncr1, Fcrl5, S1pr3, Il2*). Principal Component Analysis (PCA) based on the gene expression data showed a lack of independent clustering among these experimental conditions with PC1 and PC2 accounting only for 22% and 9% of variance, respectively (Figure 5A). Pearson correlation comparisons among the different samples also indicated that gene expression was similar across all experimental groups (Score of 0.99). In contrast to gene expression profiles, PCA analysis of genome wide OCRs highlighted significant differences between the groups, which were largely driven by the strength of TCR signaling (N4 versus T4), as shown in PC1 (Figure 5B and Figure S6A). Lowering IL-2 signaling also contributed to differential chromatin remodeling that was most pronounced when OT-I cells received strong (N4) compared to weak (T4) TCR signals (in PC2). This was further confirmed using Pearson correlation scores that also revealed the highest similarities in OCRs between T4- (weak) primed OT-I memory cells, independent of IL-2 signaling strength, and the lowest similarities between N4- and T4-primed OT-I memory cells. In addition, Pearson correlation comparisons of OCR profiles among the various experimental conditions indicated that chromatin landscapes were substantially different (score of 0.75). Collectively, these results suggested that the strength of TCR and IL-2 signaling orchestrates memory CD8^+^ T cell functional programming through the modification of chromatin accessibility but not *de novo* gene expression.

**Figure 5.**
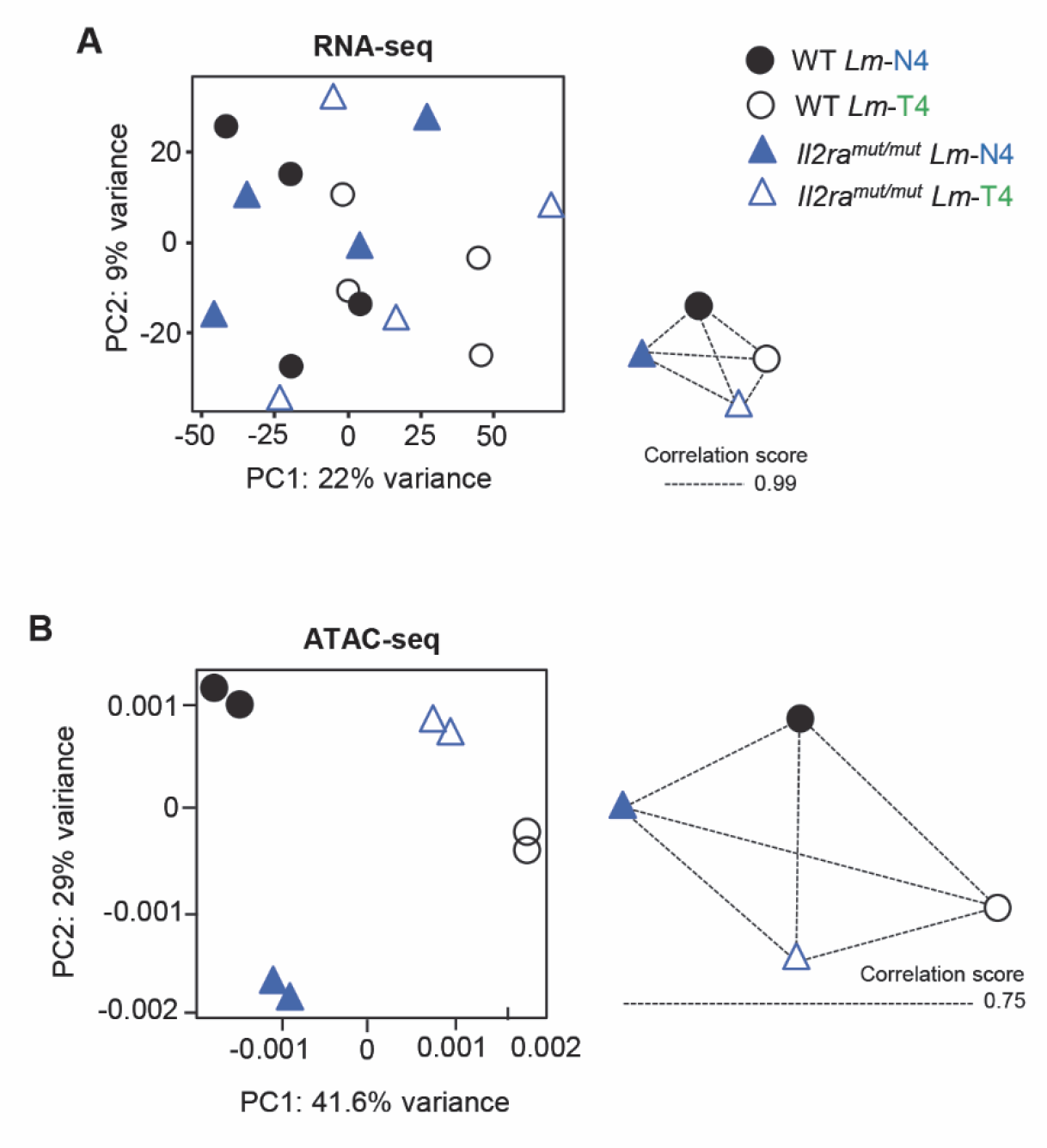
TCR and IL-2 signals alter chromatin remodeling but not gene expression in memory CD8^+^ T cells. *Il2ra^mut/mut^* and WT OT-I cells were adoptively transferred to WT recipient mice, and infected the next day with *Lm*-Ova N4 or T4. 4 weeks later, OT-I memory cells were sorted by FACS and processed for RNA-seq and ATAC-seq analysis. Principal component analysis (PCA) of **(A)** RNA-seq and **(B)** ATAC-seq datasets. Correlation network of similarity between each condition in gene expression (RNA-seq) or chromatin accessibility (ATAC-seq) is also shown. Edge length corresponds to similarity (Pearson correlation). RNA-seq (n=4 mice) and ATAC-seq (n=3-4 mice) results were summarized from 2 independent replicates of experiments.

### Modulating both TCR and IL-2 signaling strength together is required to induce broad chromatin remodeling

To determine whether the modulation of TCR, IL-2 or both signals induce distinct, overlapping or synergistic sets of OCRs genome-wide, we first conducted a side-by-side comparative analysis of the unique OCRs between *Il2ra^mut/mut^* and WT OT-I memory cells primed with *Lm*-expressing strong (N4) versus weak (T4) APLs (comparison 1, Figure 6A). Distribution of OCRs across the genome suggests a greater enrichment for TSS regions (<1kb) in OT-I memory cells induced upon T4 versus N4 priming (Figure S6B). This first level comparison enabled to identify the OCRs that are unique when i) TCR signaling strength varies independent of IL-2 signals (A and B) and ii) when IL-2 signaling strength varies independent of TCR signals (X and Y). Next, through a second level comparison of the unique OCRs revealed by the first comparisons (A versus B and X versus Y), we defined which OCRs were induced by modulating either i) TCR but not IL-2 signaling strength, ii) IL-2 but not TCR signaling strength, or iii) both TCR and IL-2 signaling strength (comparison 2, Figure 6A). This analysis revealed 6,919 unique OCRs (∼41%), i.e., 4,852 potentially associated genes (defined as the nearest, see Methods), common between *Il2ra^mut/mut^* and WT memory cells that received weak versus strong TCR priming signals; therefore, these OCRs were induced upon modulating the strength of TCR signaling whether or not intact IL-2 signals were present (Figure 6B). Only 722 OCRs (∼6%), i.e., 663 potentially associated genes, that were common to the memory cells that received either strong or weak TCR signals, were uniquely induced by altering IL-2 signals, establishing that modulating the strength of each signal orchestrates the accessibility of different regions of the chromatin. Most interestingly, when both TCR and IL-2 signaling strengths were changed together, this led to the opening of a completely new set of 12,406 or 12,249 unique OCRs, i.e., respectively 7,174 or 7,124 potentially associated genes, depending on the comparison run. As expected, the unique OCR-associated genes in both comparisons almost fully overlapped (98%, Figure S6C). This result suggested that intact IL-2 signaling alone could not rescue weak TCR priming signals to remodel chromatin, but rather acted synergistically with TCR signals during priming to program chromatin accessibility in memory cells.

**Figure 6.**
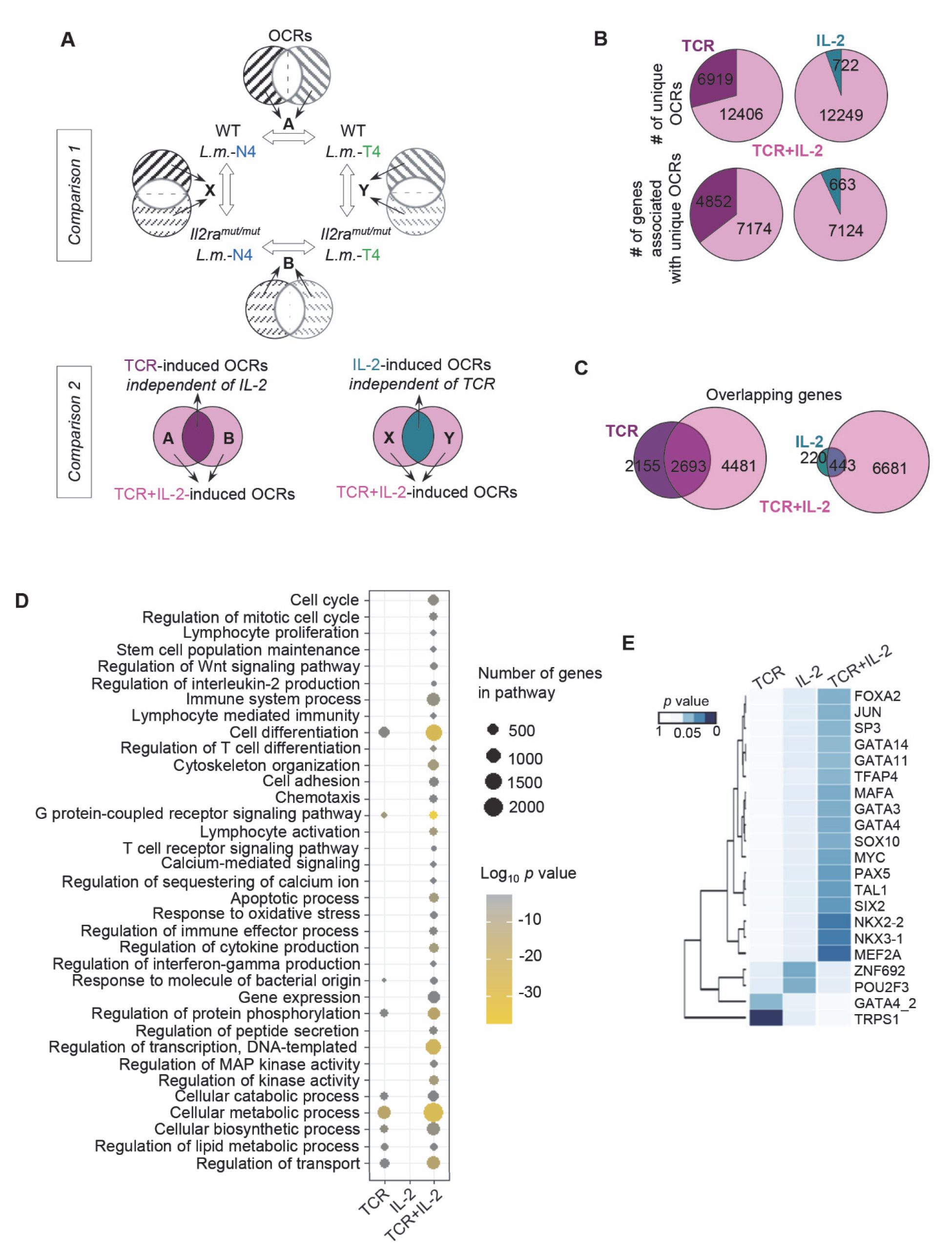
Extensive chromatin remodeling in memory CD8^+^ T cells is observed upon combined changes in both TCR and IL-2 signaling strength. **A** Schematic of the comparative analyses conducted between the open chromatin regions (OCRs) in OT-I memory cells (WT, *Il2ra^mut/mut^*) primed following *Lm*-N4 versus *Lm-*T4 infection. **(B)** Pie charts show the number of unique OCRs and related genes with unique peaks controlled by changes in TCR, IL- 2 or TCR+IL-2 signaling from comparisons depicted in (A). **(C)** Venn diagrams show the number of unique or overlapping genes associated with peaks induced by modulating TCR, IL-2 or TCR+IL-2 signals. **(D)** Network analysis of biological-process GO term enrichment of the genes with unique peaks induced by changes in TCR, IL-2 or TCR+IL-2 signals. Over- represented GO terms were analyzed by Panther and visualized in a scatterplot. Node color is proportional to the FDR-adjusted *p*-value of the enrichment. Node size is proportional to the number of genes enriched in each GO term. **(E)** Heat map shows the transcription factors enriched in unique OCRs induced by modulating TCR, IL-2 or TCR+IL-2 signals. ATAC-seq (n=3-4 mice) results were summarized from 2 independent replicates of experiments.

Of the genes associated with unique OCRs induced by changes in TCR or IL-2 signaling strength, a substantial proportion of them (∼55% to 67%) overlapped with the genes associated with unique OCRs induced by modulating both TCR+IL-2 signals (Figure 6C and Table S3). However, more than 60% of the genes associated with OCRs induced by modulating TCR+IL-2 signals were unique and did not overlap with the genes with unique OCRs induced by either varying TCR or IL-2 signaling strength only. Biological process gene ontology (BP-GO) analysis of the non-overlapping genes from Figure 6C revealed significantly greater diversity of biological processes regulated by modulating both TCR+IL-2 signals compared to TCR only while changes in IL-2 signaling did not yield any significant GO pathways (Figure 6D, Table S4). The pathways associated with the genes exhibiting unique OCRs made accessible by TCR+IL-2 signal changes were related to cell cycle, proliferation and IL-2, stem cells, T cell activation, differentiation and immunity, G-protein-coupled receptor and calcium signal transduction, transcription, chemotaxis and adhesion, cytokine response, apoptosis, metabolism and catabolism. In contrast, the pathways associated with OCRs induced by changes in TCR signaling strength were much more restricted and included immune defense, G-protein coupled receptor and metabolic/catabolic biosynthetic processes.

We next hypothesized that specific groups of transcription factors (TFs) may bind more selectively to the distinct OCRs and contribute to drive specific functional features of the memory cells. Using a motif analysis approach, we then identified candidate TF-binding motifs across OCRs induced upon modulating TCR, IL-2 and TCR+IL-2 signals respectively, and performed enrichment analysis for known motifs based on the HOMER/JASPAR database. We next ran a PCA analysis to determine which TFs may account for segregating the three sets of OCRs (Figure 6E). We revealed 21 TFs, among which 17 that were enriched in the OCRs induced by changes in TCR+IL-2 signals, while only two were enriched in OCRs induced by varying TCR or IL-2 signals only. Altogether, these results supported the idea that modulating the strength of both TCR and IL-2 signals together programmed memory CD8^+^ T cells with a greater set of OCRs and putative TFs binding sites, consistent with an improved functional fitness.

### Genes involved in memory CD8^+^ T cell stemness, cell cycle and calcium fluxes have promoter regions accessible to key transcription factors

To further link the reported changes in global chromatin accessibility to functionally relevant genes in memory CD8^+^ T cells, we next focused on the genes that exhibited differentially accessible OCRs in the promoter area close to the TSS (+/-3 kb) in WT and *Il2ra^mut/mut^* OT-I memory cells primed with weak or strong TCR signals. We postulated that such OCRs were more likely to reflect direct phenotypic and functional differences in memory CD8^+^ T cells. We noted differentially accessible OCRs in clusters of genes related to stemness, cell cycle and calcium fluxes, which all represent important hallmarks of T cell functional fitness and activation (Figures 7A and Table S5). The greater chromatin accessibility in the promoters of most of these genes in OT-I memory cells primed with strong TCR and IL-2 signals (i.e., WT, *Lm*-N4) was remarkable while these regions were largely closed in those that received weak TCR and IL-2 signals (i.e., *Il2ra^mut/mut^, Lm*-T4). In T cells that received either weak TCR or IL-2 signals, chromatin accessibility was more nuanced (even if TCR signals had a stronger impact), suggesting that other mechanisms also contributed to their accessibility.

**Figure 7.**
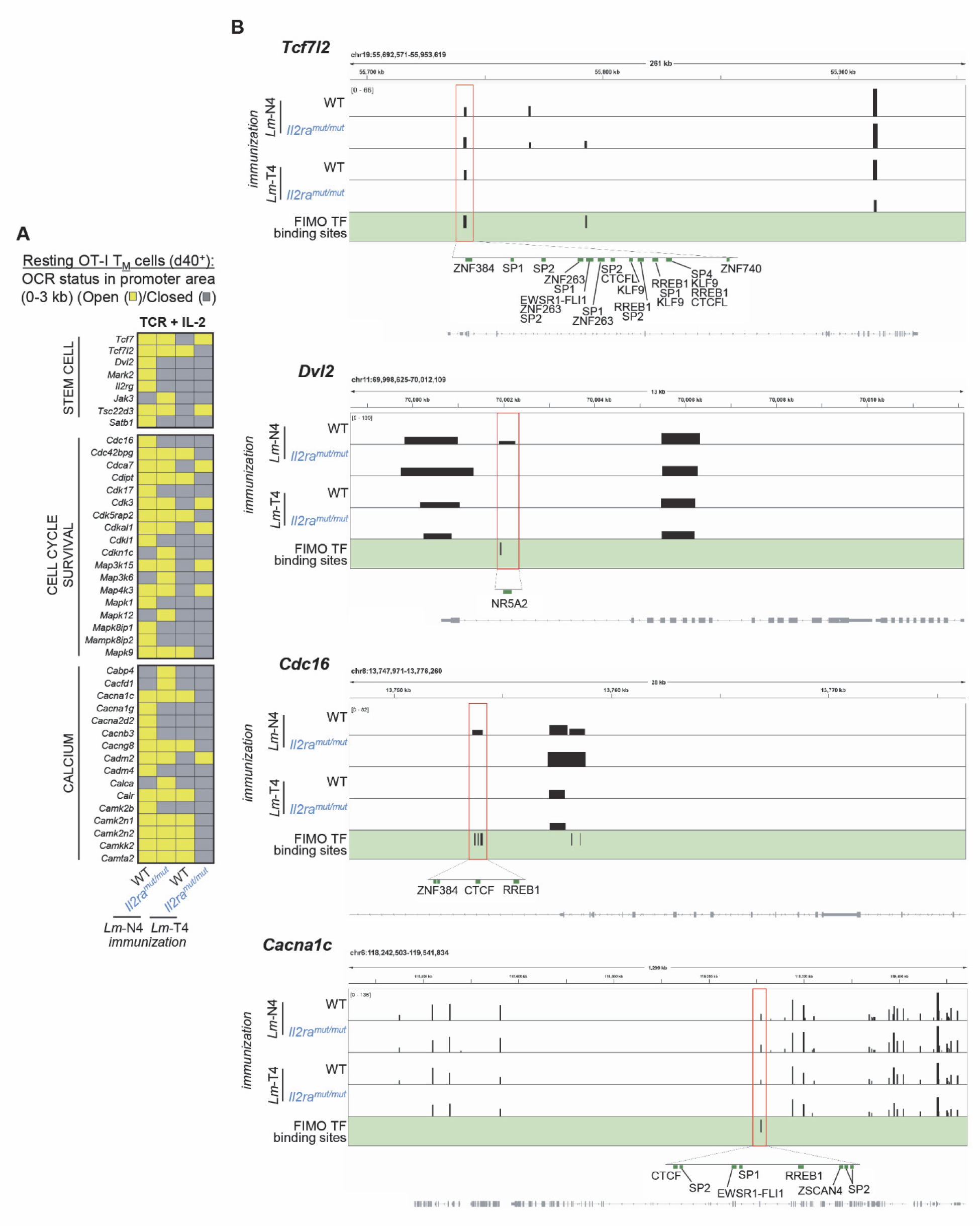
Chromatin accessibility and predicted TF binding in promoter areas of stem cell, cell cycle and calcium flux-related genes of memory CD8^+^ T cells primed under distinct strengths of TCR and IL-2 signals. (A) **Heat map shows unique OCRs (yellow) in genes** encoding stem cell-related proteins, cell cycle/proliferation and calcium fluxes that were revealed in Figure 6B when TCR and IL-2 signals were modulated together, in the promoter area and close to the TSS (+/- ∼3kb). Chromatin region status (open, yellow/closed, grey) in WT and *Il2ra^mut/mut^* OT-I memory cells primed with *Lm*-N4 or *Lm*-T4 are reported. **(B)** Selected examples of genes (*Tcfl2, Dvl2, Cdc16, Cacnac1*) with differential OCRs (red box) in these OT-I memory CD8^+^ T cells. Higher magnifications of the OCRs show FIMO-predicted binding sites for indicated TFs.

We next selected genes involved in the distinct biological processes highlighted above that included *Tcf7, Tcf7l2, Dvl2, Mark2*, *Il2rg, Cdc16, Cdk17, Mapk9, Cacna1c, Cacng8. Tcf7, Tcf7l2, Dvl2* and *Mark2* are involved in the activation the Wnt/β-Catenin pathway while IL-2Rγ transduces common γ-chain cytokine signals (i.e., IL-2, IL-7 and IL-15). *Cdc16*, *cdk17* and *Mapk9* products control cell cycle progression whereas *Cacna1c* and *Cacng8* encoding proteins cooperate to form and regulate L-type high voltage calcium channels implicated in intracellular calcium accumulation. We then evaluated likelihood of binding for known TFs in the differentially accessible promoter-region OCRs of these genes using Find Individual Motif Occurrence (FIMO ; Figure 7B and S7B and Table S5). While we did not find any TFs binding sites in *Tcf7, Mark2* or *Cdk7* promoter OCRs, we revealed binding sites for multiple TFs in *Tcf7l2, Dvl2, Il2rg, Cdc16, Mapk9, Cacna1c* and *Cacng8* promoter OCRs.

Notably, OCRs associated with the promoter regions of selected genes, such as *Tcfl2*, *Cdc16*, *Cacna1c*, *Cacng8* and *Il2rg*, contained a greater number of putative TF binding sites and higher diversity in families of TFs (i.e., SP, ZNF, KLF, CTCF, RREB) that could potentially bind at these sites, suggesting these OCRs and TFs could play important roles in regulating expression of these genes (Figure 7B and Table S5). In contrast, other OCRs in genes like *Dvl2, Cdkl1* and *Mapk9* only exhibited limited TF binding sites, suggesting that these areas may either be less important or more specific in regulating expression of these genes. Thus, this analysis revealed that many of the differentially accessible OCRs in genes potentially involved in stemness, cell cycle and calcium fluxes may be subjected to further regulation by families of TFs, consistent with a greater functional fitness of the memory cells primed by strong TCR and IL-2 signals.

### The greater chromatin accessibility in memory CD8^+^ T cells primed with strong TCR and IL-2 signals is consistent with their higher subset diversity and phenotypes

If the epigenetic differences observed in both the breadth of OCR-related GO pathways and increased accessibility of stem-cell related gene promoters in memory CD8^+^ T cells induced by strong TCR and IL-2 signals reflects functional relevance, memory CD8^+^ T cells primed with strong signals should give rise to a greater diversity of subsets and phenotype than those that received weak TCR and/or IL-2 signals. Using high dimensional flow cytometry with a 26-color panel that included cell-surface markers, intracellular functional markers and TFs relevant to CD8^+^ T cells (Table S6), we conducted an in depth characterization of resting OT-I memory cells primed with weak or strong TCR and IL-2 signals (Figure 8 and Figure S8). Focusing first on known memory CD8^+^ T cells subsets such as terminally differentiated KLRG1^+^ or CX3CR1^+^CD27^-^ effector memory (TEM) cells and CD127^+^KLRG1^-^ or CX3CR1^-^CD27^+^ central memory (TCM) cells ^14, 30^ revealed a higher proportion of TEM-like cells when primed with intact versus weak IL-2 signals (Figure 8A), a finding consistent with IL-2 signals promoting differentiation of robust effector cells ^16, 31^. Interestingly, however, increasing TCR signals neither altered the proportion of TEM or TCM cells, underscoring the need to achieve deeper resolution of memory subsets to reveal more granular differences.

**Figure 8.**
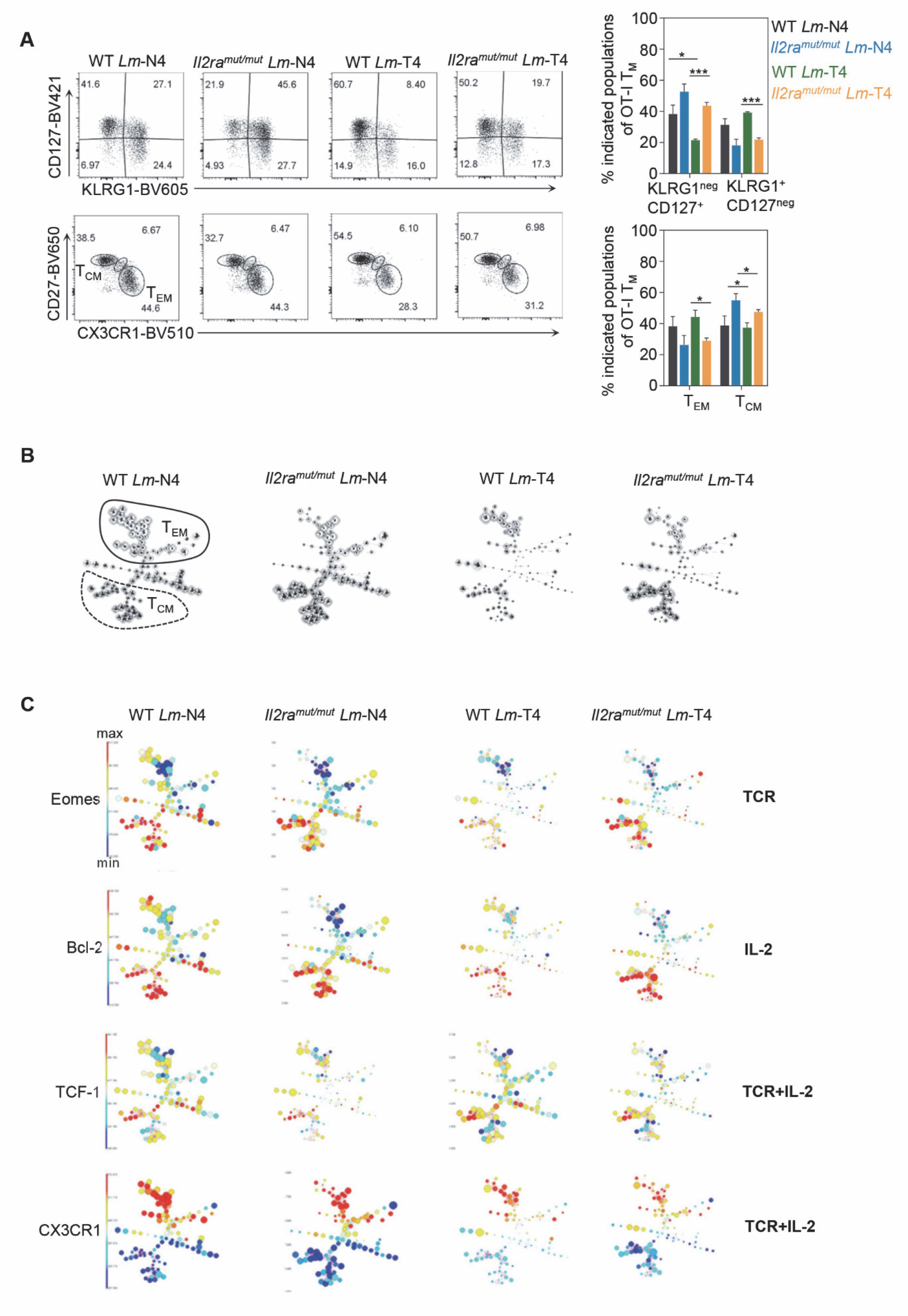
Robust TCR and IL-2 signals induces the most memory CD8^+^ T cell subset diversity and phenotypes. **A** FACS analysis of *Il2ra^mut/mut^* and WT OT-I memory cells primed after infection either *Lm*-N4 or *Lm*-T4 ∼5 weeks later, after staining with a panel of 26 memory cell-relevant markers. Classically defined effector (TEM), central memory (TCM) subsets or based on cell-surface expression of KLRG1, CD127, CD27 and CX3CR1 are shown. **(B)** FlowSOM analysis of each experimental group containing a pool of concatenated OT-I memory cells from 3 mice. Where classically defined TEM and TCM subsets are, is manually circled. **(C)** The expression level of indicated markers (Eomes, Bcl-2, TCF-1, CX3CR1) within each node, regulated by each or combined priming signal(s), is represented in a color scale. Data are pooled from 3-4 mice across 2 replicate experiments.

Thus, we next processed the analysis of our high dimensional flow cytometry data using flow self-organizing map (FlowSOM) that enables unsupervised clustering and dimensionality reduction, and the visualization of discrete memory cell subsets and their relative proportions (Figure 8B). This analysis revealed a remarkable heterogeneity of potential subsets in TEM and TCM cells. Memory cell subset diversity and relative proportions were closely similar in *Il2ra^mut/mut^* or WT OT-I cells (only IL-2 signals differ) primed with either *Lm*-expressing N4 or T4 epitopes. In contrast, memory cells (whether *Il2ra^mut/mut^* or WT) primed with *Lm*-expressing N4 or T4 epitopes (only TCR signaling strength differed) exhibited greater subset diversity and included many differentially represented subsets. Importantly, OT-I memory cell subset diversity and relative proportions in *Il2ra^mut/mut^* OT-I cells primed with T4 compared to WT counterpart primed with N4 or vice-versa (when both TCR and IL-2 signals differ), had the most differentially represented subsets. Deeper FlowSOM analysis on individual cell-surface and intracellular marker expression revealed the relative effect of each or combined signal(s) on the FlowSOM outlined subsets of memory cells (Figure 8C and Figure S8). In particular, markers associated with a TCM cell phenotype (TCF-1, Eomes, Bcl-2, Sca-1, CD122, CD127, CD27) exhibited significantly higher expression in the subsets included in the broad category of TCM cells. The combined modulation of both TCR+IL-2 signals at priming had the most impact on the majority of markers (e.g., TCF-1, CX3CR1 and most others), also consistent with a greater chromatin accessibility in the promoter of stem-cell related genes (e.g., *Tcf7, Tcf7l2, Il2rg*) in memory cells primed under strong TCR and intact IL-2 signals (Figure 7). Of note, some markers were more selectively affected by TCR (i.e., Eomes, IRF4) or IL-2 (i.e., Bcl-2) signal changes. Thus, collectively, these results are in agreement with our chromatin landscape analyses indicating that modulating TCR, IL-2 or TCR+IL-2 signaling strength sets distinct non-overlapping OCRs that are likely to be linked to the phenotypic output of potential memory cell subsets.

### Differentially accessible OCRs in genes involved in the control of stemness, cell cycle and calcium fluxes in memory CD8^+^ T cells correlate with distinct functional responses

As an attempt to further link the reported changes in chromatin accessibility to functional differences in the memory CD8^+^ T cells, we focused on the specific clusters of genes with differential accessibility in the promoter area revealed in Figure 7. We hypothesized that such OCRs were likely to be essential in the rapid and differential modulation of memory CD8^+^ T cell activation *in vivo*. Using FACS analysis, we tested whether WT or *Il2ra^mut/mut^* OT-I memory cells induced upon immunization with *Lm*-expressing weak (T4) or strong (N4) Ova APLs, exhibited different abilities to undergo rapid intracellular calcium signaling, enter cell cycle (16 hrs), and expand (day 4, 6, 9 and 12) after *Lm*-N4 challenge infection (Figure 8A).

Intracellular calcium accumulation early on after the secondary challenge infection (16 hrs) was significantly higher in N4-primed OT-I memory cells compared to T4-primed counterparts, whether on the WT or *Il2ra^mut/mut^* background (Figure 8B). This indicated that higher calcium signaling in memory CD8^+^ T cells largely depended on strong TCR signals at the time of priming, despite many relevant OCRs being differentially accessible when both TCR and IL-2 signals varied (Figure 7). However, we also noted that *Cabp1* (Calcium binding protein 1) and *Camk1* (Calcium/calmodulin-dependent protein kinase) accessible OCRs were only modulated by TCR signaling strength, suggesting these genes could potentially act as key regulators of intracellular calcium fluxes in early reactivated memory CD8^+^ T cells (Figure S9A).

At early stages after activation, we also found that a significantly greater proportion of T4- induced *Il2ra^mut/mut^* and WT OT-I memory cells (factor of ∼6 and 3, respectively) remained in the G0 phase while N4-counterparts had already transitioned to the G1 phase (Figures 9C and S9B). As expected, a much higher proportion of naïve T cells were in G0 compared to early activated N4-memory or effector (day 5) cells (up to ∼20 times more). In addition to their faster initiation of cell cycle, N4-induced WT OT-I memory cells underwent more robust and competitive later expansion than both WT and *Il2ra^mut/mut^* OT-I memory cells primed with the weak T4 agonist (Figures 9D and S9C). Memory cell numbers and frequencies quantified 6, 9, and 12 days post- *Lm*-N4 challenge were significantly lower (factor of 2-4) when OT-I cells had been primed with weak TCR and IL-2 signals. These results showed that rapid memory cell entry into cell cycle and competitive proliferation/expansion largely depended on the combination of robust TCR and IL-2 signals during priming. These findings were also consistent with the differentially accessible OCRs noted in the promoters of cell-cycle checkpoint (i.e., *cdc16, cdkl1, cdkn1c*), stem-cell (*Tcf7l2, Dvl2*) and IL-2 (*Il2rg, Jak3, Tsc22d3*) encoding genes (Figure 7).

**Figure 9.**
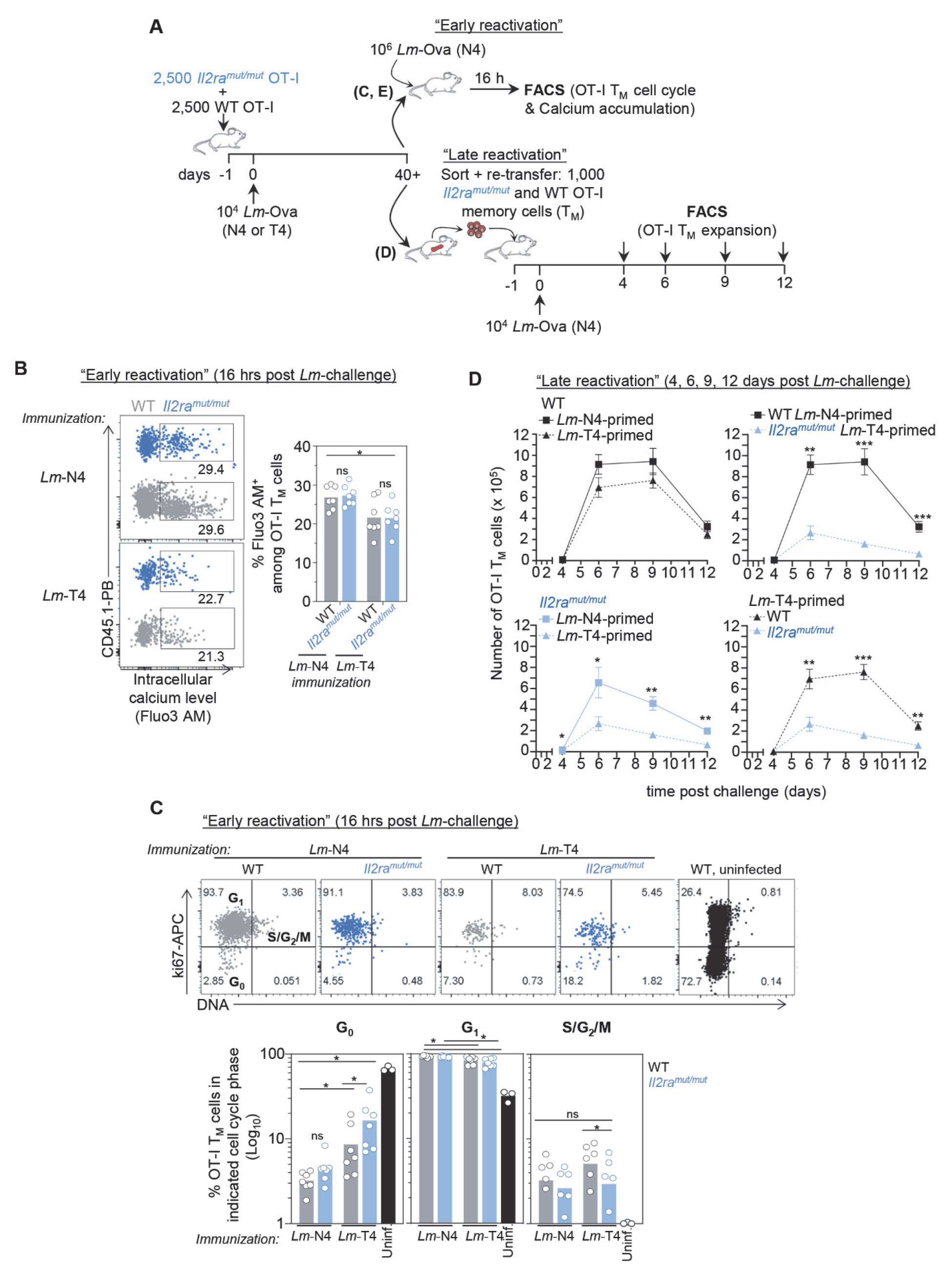
Memory CD8^+^ T cells induced with robust TCR and IL-2 signals initiate faster calcium signaling, cell cycling and expand more competitively than those primed with weaker signals. **A** Schematic of experimental designs used in (B-D). Briefly, 2,500 *Il2ra^mut/mut^* or WT OT-I Td^+^ cells were adoptively transferred to WT naïve hosts, and infected the next day with *Lm*-Ova N4 or *Lm*-Ova T4, and housed for ∼40 days before the next steps. In **(B, C)**, immunized mice were challenged with 10^6^ *Lm*-Ova N4 and 16 hrs later (“early activation”), spleen cells were either (B) loaded with Fluo3 AM or not (C) and stained for cell-surface CD3, CD8, congenic markers. In (C) cells were stained for intracellular expression of Ki67 and DNA. Representative dot plots comparing *Lm*-OvaN4 versus *Lm*-Ova T4-primed reactivated OT-I memory cells for indicated marker expression. Graphs summarize the average of 3-4 mice for the 4 experimental conditions. In **(D)**, 1,000 *Il2ra^mut/mut^* and WT OT-I memory cells from each priming condition, were FACS-sorted and transferred at a 1:1 ratio to new hosts subsequently infected with *Lm*-OvaN4 the day after. At indicated days, spleen cells were stained to quantify *Il2ra^mut/mut^* and WT OT-I memory cell expansion and numbers. Data are the pool of 2 independent replicate experiments (n=8-14mice). *p*-values are indicated when relevant with \**p* < 0.05; \*\**p*< 0.01; \*\*\**p*< 0.001; \*\*\*\**p*< 0.0001; NS, not significant, using two-tailed unpaired Student’s *t* test.

In summary, differential chromatin accessibility at promoter area of genes related to intracellular calcium processes, cell cycle, proliferation and stemness, correlated with distinct functional responses of reactivated memory cells induced by strong or weak TCR and/or IL-2 signals. Consistent with our epigenetic analysis, these results support a model of greater fitness of memory cells primed with stronger TCR and IL-2 signals while those that received weak TCR and/or IL-2 signals were sluggish to rapidly activate, enter cell cycle and expand.

## Discussion

Here, we reveal that the strength of TCR signaling is a key driver of memory CD8^+^ T cell functional characteristics and fitness. We establish that the amount of TCR signals integrated by naïve CD8^+^ T cells during priming determines whether IL-2 signals are required for them to form fully functional memory CD8^+^ T cells, i.e., that are able to undergo rapid reactivation (calcium signaling, cell cycling) and competitively expand during a rechallenge infection, and that exhibit higher stemness characteristics. We also show that the concomitant modulation of TCR and IL-2 signals has a profound impact on memory CD8^+^ T cell epigenetic landscape, as measured by a significantly increased chromatin accessibility. In contrast, the extent of chromatin remodeling and opening in memory cells primed under lower TCR or IL-2 signaling strengths was much more reduced.

Current dogma states that increasing the strength of TCR signals in CD8^+^ T cells i) promotes an effector versus a memory CD8^+^ T cell fate ^10, 11^, ii) augments the size of primary CD8^+^ T cell expansion and of the memory cell pool ^7, 19^, and iii) does not alter the functional features of memory cells ^19^. While our results confirm prior findings (i) and (ii), they nevertheless challenge the concept (iii) that TCR signaling strength does not modulate the functional attributes and programming of memory CD8^+^ T cells. Although CD8^+^ T cells primed by weak TCR signals give rise to significantly lower numbers of memory cells most likely as a result of curtailed expansion ^19^, our data also show i) that lowering TCR signaling strength alters their programming through distinct chromatin remodeling and ii) the requirement for intact IL-2 signaling in order to form fully functional memory cells.

The importance of IL-2 signals in enabling the formation of functional memory cells has been previously documented ^32^, but it has also remained controversial ^33^. We confirm here the results of the prior studies by showing that T cells primed with the natural OvaN4 epitope do not require IL-2 signals during priming ^33^ whereas those primed with the LCMV-derived GP33-41 epitope do ^32^. By establishing that TCR signaling strength, whether as a result of weak epitope/MHC stability or epitope/MHC/TCR interactions, is the major variable controlling CD8^+^ T cell-dependency on IL-2 signals during priming, we further provide a rational explanation for the earlier discrepancy. Importantly, these results are also consistent with the idea that a quantum threshold of TCR signals needs to be reached for naïve CD8^+^ T cells to bypass the need for intact IL-2 signals and form a pool of functional memory cells. Below such threshold, even though CD8^+^ T cells still expand and form memory, they fail to become fully functional, e.g., equipped to undergo optimal reactivation, initiate cell cycle and competitively expand. While this may represent a safeguard mechanism to prevent memory T cell clones to be reactivated against self- or tumor-derived antigens of low affinity ^20^, this result also suggested that the induction of effective memory CD8^+^ T cell responses against weak T cell epitopes, requires robust IL-2 signaling. These findings are particularly relevant in the context of recent studies using an IL-2 “superkine” partial agonist that preferentially triggers the ERK pathway in spite of STAT5 downstream of the IL-2 receptor, and skews CD8^+^ T cell differentiation towards a stem cell fate ^34, 35^. Our results suggest that the acquisition of such superior functional features through IL-2 signaling is very likely to be also regulated by the amount of TCR signals received by naïve CD8^+^ T cells during priming. These findings have important implications for improving the rationale design of vaccines and T cell therapies targeting low affinity tumor-derived antigens in particular.

Approximately one-third of the naturally selected microbial pathogen-derived epitopes we have studied appear to induce CD8^+^ T cells that are dependent on intact IL-2 signaling to differentiate into memory cells that competitively re-expand during antigen re-encounter. This suggests that this mechanism is likely to extend beyond the Ova model system, increasing its potential significance in epitope selection and vaccine design. While our data are consistent with such possibility, it has also been established that Foxp3^+^ regulatory T (Treg) cells favor the priming of high versus low affinity polyclonal CD8^+^ T cells ^36^. It is therefore conceivable that IL-2 produced in close proximity to low affinity naïve CD8^+^ T cells that cannot effectively bind IL-2 (i.e., *Il2ra^mut/mut^* or *Il2ra^-/-^*), is consumed by Treg cells, further promoting primary expansion of higher affinity clones with a distinct TCR repertoire that would require IL-2 signals during priming for competitive re-expansion. Whether Treg cells are involved or not in expanding a distinct set of T cells, our data together with current literature, remain consistent with the concept that IL-2 plays an essential role in the functional imprinting of memory CD8^+^ T cells.

The findings presented in this work are also consistent with several studies showing that the strength of TCR signaling in naive CD4^+^ T cells is an essential determinant of T helper cell effector fates, for which strong TCR signaling skews their differentiation towards Th1 ^37^, follicular helper (TFH) ^38^ or germinal center (GC-TFH) CD4^+^ T cells ^39^. TCR/peptide MHC dwell times rather than equilibrium binding was proposed to predict T helper effector cell fates, with long dwell times promoting TFH or GC-TFH rather than Th1 effector cell fates ^39^. Recent reports further involved IL-2 and its signaling as an important mechanism orchestrating such CD4^+^ T cell fates ^40, 41^. In one study, strong TCR-stimulation with long dwell times skewed naïve CD4^+^ T cell differentiation into effector TFH cells (Bcl-6^+^) and endowed them with IL-2-producing capacity, consistent with a model in which IL-2-secreting cells are precursors of TFH cells ^41^.

Non-IL-2-secreting CD4^+^ T cells, that received weaker TCR signals and IL-2 from IL-2- secreting CD4^+^ T cells, rather differentiated into non-TFH effector cells (Blimp-1^+^). In contrast, in the other study, strong TCR stimulation, which induces high levels of cell-surface CD25, skewed CD4^+^ T cell-differentiation into Th1 effector cells, while weaker TCR stimulation, and therefore low levels of cell-surface CD25, was associated with a Th1 and TFH memory cell fate ^40^. In line with our results and the literature on CD8^+^ T cells ^10, 19^, this latter study showed that decreasing TCR signals induced memory CD4^+^ T cells that competitively expanded and produced effector functions. Whether weakly stimulated CD4^+^ T cells lacking IL-2 or even other cytokine signaling pathways would fail to form functional memory cells is not known and would need further investigations.

Another key message of this work is that the combination of strong TCR and IL-2 signals during CD8^+^ T cell priming together imposes important modifications in the chromatin accessibility of memory CD8^+^ T cells, therefore alters naïve CD8^+^ T cell long-term programming. We report that combining these signals enables the opening of a distinct genome-wide epigenetic landscape that only moderately overlaps with the landscapes modulated by either TCR or IL-2 signals and appears much broader. The breadth of biological processes targeted and the greater number of TFs (17 compared to 2) that can potentially bind to accessible chromatin regions uniquely present in memory cells primed by strong TCR and IL-2 signals, support the interpretation that combining these signals together induces synergistic mechanisms of chromatin remodeling that are not turned on by each individual signal. It is likely that establishment of these distinct epigenetic landscapes occurs at early effector stages after priming and involve DNA/histone-modifying methyltransferases that control chromatin accessibility in memory precursor or effector cells ^23, 24^. Another non-exclusive possibility based on our data is that combinations of TCR- and IL-2-dependent transcriptional regulators could act together as genome organizers. Prior reports have shown that IRF4 and BATF TFs for instance bind DNA cooperatively to determine initial chromatin accessibility and initiate specific STAT-dependent transcriptional programs in Th17 CD4^+^ T cells ^42^ and early effector CD8^+^ T cells ^43^. Our transcriptomic analysis at early stages post-priming highlighted that genes encoding both IRF4 and BATF3 TFs were upregulated in OT-I cells primed with robust TCR signals, which could contribute to establishing the distinct epigenetic landscape that we observed.

The analysis of the differentially accessible OCRs in the promoter areas of resting memory cells primed with robust TCR and IL-2 signals revealed clusters of genes involved in stem-cell features, cell cycling and calcium fluxes. Our FIMO analysis further indicated that many TFs belonging to distinct families, can potentially bind to these areas. This finding suggested that these differentially accessible promoter regions may play important roles in the expression of these various genes and the functional features of the memory cells. Both the higher chromatin accessibility in genes promoters associated with the Wnt/β-catenin pathway as well as the genome-wide enrichment in predicted binding sites of TFs such as GATA-3, TAL-1, NKX3-1, SIX2 which act as lineage commitment TFs and are involved in stem cell maintenance ^44, 45^, are consistent with the superior functional features and subset diversity of the memory cells primed under robust TCR and IL-2 signals.

Our comparative high dimensional FACS analysis of resting memory CD8^+^ T cells is also consistent with differences in stem-cell related gene promoter accessibility. While subdividing the memory cells based on TEM and TCM subsets failed to reveal differences other than those likely accounted for by intact IL-2 signaling (which favors KLRG1^+^ effector cell differentiation ^16, 31^), FlowSOM analysis and visualization achieved deeper resolution of potential subsets of memory cells. This enabled a more precise assessment of how much the strength of each signal modulated the size of individual subsets and expression levels of a given memory cell marker. This highlighted that when both TCR and IL-2 signals were co-modulated, the diversity of subsets and their expression of markers driving stemness (TCF-1, Eomes, Sca-1, CD127, CD62L, CD44) were clearly increased. Importantly, the better chromatin accessibility in promoters of genes controlling cell cycle initiation, calcium fluxes and proliferation in resting memory CD8^+^ T cells primed with robust TCR and IL-2 signals correlated with the functional outcomes after their reactivation. These memory cells accumulated intracellular calcium, initiated cell cycle and expanded significantly more than those that were primed with low TCR and IL-2 signals. Since multiple TFs were predicted to bind in these OCRs, it is conceivable that several levels of regulation are expected to take place. Interestingly, providing IL-2 signals to the IL-2 signaling defective memory CD8^+^ T cells rescues their ability to competitively expand, indicating that optimal clonal expansion can be achieved through STAT5 signaling. However, given the genome wide differences in chromatin accessibility between the memory cells primed with distinct TCR and IL-2 signals, it seems unlikely that giving IL-2 signals during reactivation of the memory cells primed with weak TCR and IL-2 signals will be sufficient to rescue an epigenetic landscape associated with “high fitness”.

## METHODS

### Ethics Statement

This study was carried out in strict accordance with the recommendations by the animal use committee at the Albert Einstein College of Medicine. All efforts were made to minimize suffering and provide humane treatment to the animals included in the study.

### Mice

All mice were bred in our SPF animal facility at the Albert Einstein College of Medicine. We used wild-type (WT) C57BL/6J (B6) 6-8 weeks old male or female mice, congenic CD45.1^+/+^ (JAX#002014), OT-I^+^ (JAX#003831), P14 (JAX#004694, backcrossed to B6/DBA/2>6 times), CD11c-DTR^+/-^ (JAX#004509) and Rosa26-Actin-tomato-stop^loxP/loxP^-GFP (TdT)(JAX#007576) all purchased from the Jackson labs. *Il15^-/-^* (stock#4269) mice were purchased from Taconic farms. We also bred gBT-I ^46^ (gift Dr. Carbone), L9.6^+^ Kd^+^^47^, *Il2ra^mut/mut^* ^28^ and *Ifnar^-/-^* mice (gift Dr. Kohlmeier, Emory Vaccine Center). All mice are on the B6 genetic background unless otherwise specified.

### Microbial pathogens and mouse infections for primary and memory response analyses

#### Listeria monocytogenes (Lm) bacteria and infections

*Lm* on the 10403s genetic background ^48^ was used to express different antigens: Ovalbumin (*Lm*-Ova257-264 (N4)) and its APLs (T4, A8, Q4), Herpes Simplex Virus 2 (*HSV-2*) glycoprotein B (*Lm*-gB498-505), lymphocytic choriomeningitis virus (*LCMV*) glycoprotein (*Lm*-GP33-41, *Lm*-GP276-286). *Lm*-expressing N4, T4, Q4, GP33-41, gB498-505, GP276-286 were obtained from D. Zehn ^19^. We generated *Lm*-A8 according to published methods ^49^. Briefly, a DNA fragment encoding for a chimeric protein Ova209-309- gB447-550 containing the Ova A264 (A8) mutation and the gB498-505 epitope was synthetized (Genewiz) and cloned into the pHSLV *Lm* transfer vector for *Lm* selection and expression under the LLO/*Hly* promoter. All *Lm* were prepared after passaging into WT B6 mice, by growing to log phase (OD600∼0.3-0.4) and kept as frozen aliquots for single use in -80°C. For infections, bacteria were grown to a logarithmic phase (OD600∼0.05-0.15) in broth heart infusion medium, diluted in PBS to infecting concentration (10^4^/mouse) and injected i.v. Secondary challenge infections were performed 4-6 weeks later with 10^6^ *Lm*-Ova (N4) unless otherwise specified in Figure legends.

#### Herpes Simplex virus 2 (HSV-2) strains and infections

Virus stocks, both WT HSV-2 (strain 186) and TK^-^ HSV-2 (186ΔKpn) were prepared in Vero cells using standard procedures and virus stocks were kept at -80°C. *Prior to infections, female* mice were treated with 2 mg medroxyprogesterone acetate subcutaneously (s.c.) and 5 days later were inoculated intravaginally with 2x10^5^ plaque forming units (PFU) of TK^-^ HSV-2. For primary infection, spleens were harvested 7.5 days later for FACS analysis. The WT HSV-2 virus was used for i.v. rechallenge infections 60 days after immunization and spleens were harvested 6.5 days later for FACS analysis.

#### Vesicular Stomatitis Virus (VSV) strains and infections

*VSV* encoding *Lm* listeriolysin O (LLO91-99) and p60217-225 ^50^ or Ova (gift Kamal Khanna, NYU) kept at -80°C were thawed and diluted in cold PBS right before infections. 2x10^5^ PFU/mouse were injected i.v. into mice. For primary infection, spleens were harvested 7.5 days later for FACS analysis. For rechallenge infections, mice were infected i.v. with *VSV*-LLO91-99-p60217-225 at least 40 days after immunization and spleens were harvested 6.5 days later for FACS analysis.

#### *In vitro* T cell assays

##### Antigen presenting cell (APC) preparation

Single cell suspension was prepared from WT B6 spleens, lysed with ACK lysis buffer, and incubated with 1mL of anti-CFTR in 1X PBS for 1 min at 37°C before quenching with 10 mL of complete RPMI media. The APCs were then mixed with indicated concentrations of Ova agonist peptides (N4, T4, A8) and E1 antagonist control for 2 hrs at 37°C. APCs were then rinsed and resuspended in complete RPMI media.

##### T cell preparation and co-culture setup

Single cell suspension was prepared from WT or *Il2ra^mut/mut^* OT-I spleens, lysed with ACK lysis buffer, and stained with CTV (Invitrogen) according to the manufacturer’s protocol. The APCs loaded with Ova peptides were co-cultured with CTV-labelled T cells at a 3:1 ratio in a tissue treated 96-well v-bottom plates Plates were loaded into a 37°C incubator attached to custom Tecan Freedom EVO 75 robotic platform programmed to conduct automated robotic time series that collected supernatant for cytokine analysis and cell pellets for surface/intracellular marker analysis. For cytokine quantifications, co-culture supernatants were thawed at room temperature for 1 hr and cytokines stained with mouse Th1/Th2/Th17 cytokine kit (BD) according to the manufacturer’s protocol. For proliferation and activation, cell pellets were incubated with fluorescently tagged antibodies (see Table S5) at room temperature for 30 min. All samples were next acquired on a BD Fortessa FACS HTS.

##### Preparation of cell suspensions and staining for flow cytometry analysis

Spleens or lymph nodes (inguinal and cervical) were dissociated on a nylon mesh. Cell suspensions were treated with red blood cells (RBC) lysis buffer (0.83% NH4Cl vol/vol). Blood was harvested into heparin tubes and RBC lysed. Cell suspensions were incubated with 2.4G2 Fc Block and stained with fluorescently tagged antibodies (See Table S6) in FACS buffer (PBS, 1%FCS, 2mM EDTA, 0.02% sodium azide). Brilliant stain buffer (BD) was used when more than two Abs were conjugated with BD Horizon Brilliant fluorescent polymer dyes. Ova257-264/K^b^, gB498-505/K^b^, GP33-41/D^b^, GP276-284/D^b^, LLO91-99/K^d^ and p60217-225/K^d^ biotinylated monomers (1mg/mL) obtained from the NIH tetramer Core Facility, were conjugated with PE-labeled Streptavidin (1mg/mL) as follow: 6.4 μL of PE-Streptavidin were added to 10 μL of monomers every 15 min 4 times on ice. Newly generated tetramers (1/400-1/500 dilution) were used to stain cells for 1 hour at 4C. For transcription factor (TF) intracellular staining, cells were fixed and stained according to the eBioscience Foxp3 Transcription Factor Staining Buffer Set protocol. Data acquisition was done using BD LSR II, FACSAria III or Cytek Aurora flow cytometer. All flow cytometry data were analyzed using FlowJo v9 or v10 software (TreeStar).

##### T cell sorting

CD8^+^ T cells were negatively selected from spleen using anti-CD4, anti-CD11b, anti-MHC II, anti-TER119, anti-B220 and anti-CD19 mAbs (Table S5), which all were added and incubated at 5 μg/mL for 30 min at 4C. Cells were then washed and incubated with anti-rat Ab magnetic beads at 1 bead/target cell for 40 min at 4C (Dynabeads sheep anti-rat IgG, Invitrogen). Cell suspensions were stained with Abs specific for congenic markers. Cells were sorted into 3mL of complete media (RPMI 1640, 10% FBS, 1% Penicillin/Streptomycin, 55μM β-mercaptoethanol, 1mM Sodium Pyruvate, 1X Glutamax, 1X non-essential amino acids) using a 4 laser BD FACS Aria III cell sorter. For RNA-seq experiments, cells were directly sorted into 1X lysis buffer (Takara Bio USA).

#### *In vivo* mouse assays

##### Adoptive T cell transfers

2,000 *Il2ra^mut/mut^ Cd45.1^+/+^* TdT^+^ OT-I cells and 2,000 *Cd45.2^+/+^* TdT^+^ OT-I cells from blood or spleen were prepared in PBS and transferred i.v. into *Cd45.1^+/-^* WT B6 naïve recipient mice. The next day, mice were immunized as indicated above and in Figure legends. For memory cells, 1,000 OT-I memory cells of each genotype from immunized mouse spleens, were negatively selected (see below) and flow-sorted (Aria III) based on expressed congenic markers and TdT expression. Sorted OT-I memory cells were mixed at a 1:1 ratio in 200 μl and immediately transferred to *Cd45.1^+/-^* WT B6 naïve recipient mice that were challenged the next day with indicated microbial pathogen.

##### Mixed bone marrow chimera

BM cells were obtained from flushing femurs with complete RPMI media with 10% FCS. Recipient mice were lethally irradiated with 1,200 rads before immediate reconstitution with 5x10^6^ BM cells from WT *Cd45.1^+/+^*, *Il2ra^mut/mut^ Cd45.1^+/-^* and *Il2ra^-/-^ Cd45.2^+/+^* at a 1:1:1 ratio. Mice were placed under antibiotics for 2 weeks and reconstitution ratios were checked by FACS 4-6 weeks later before immunization experiments. *IL-2/anti-IL-2 mouse rescue model:* Mice adoptively transferred with *Il2ra^mut/mut^* and WT OT-I cells were injected i.p. with a solution of 1.5μg recombinant mouse IL-2 (rIL-2, preprotech) mixed to 50μg anti-IL-2 (clone S4B6) as previously described ^32^ every day for 3 or 6 consecutive days post *Lm*-Ova infection. rIL-2 and the anti-IL-2 Ab were incubated rotating at 4°C for 1 hr prior to i.p. injection in a total volume of 300 μL of PBS per mouse.

##### In vivo blocking of antigen presentation

Mice adoptively transferred with *Il2ra^mut/mut^* and WT OT-I cells were injected with 500 *μ*g of anti-K^b^/Ova257-264 25-D1.16 mAb per mouse 2 days after immunization with *Lm*-Ova. CD11c^+^ cells were depleted in mice expressing the diphtheria toxin receptor (DTR) under the CD11c/Itgax promoter upon intraperitoneal (i.p.) injection of 4 ng/g of mouse body weight of diphtheria toxin (DT, Calbiochem) 12 hrs prior to the targeted CD11c^+^ cell depletion time as indicated in the relevant figure legends. Depletion efficiency was confirmed by FACS on 1-2 extra mice in each independent experiment and by monitoring OT-I cell expansion 7 days post immunization.

##### Cell Trace Violet (CTV) or CFSE labeling

Purified naïve OT-I, gBT, P14, or L9.6 T cells were stained with 1-5 μM of CTV or CFSE (Invitrogen) according to the manufacturer’s protocol.

5×10^4^ CTV-labeled cells were injected i.v. to congenically distinct recipient mice. Cell proliferation of CTV labeled cells was determined by flow-cytometry based on CTV fluorescence intensity dilution among cells.

##### Intracellular calcium accumulation

Splenocytes were loaded with 2.5μM Fluo3am (ThermoFisher) in PBS by incubating at 25°C for 30 min. Wells were then washed with indicator-free media and stained for flow cytometry analysis.

##### Cell cycle analysis

Following Fc-block and cell surface staining, splenocytes were fixed and stained with anti-Ki67 mAb and Live/Dead Aqua (ThermoFisher) to quantify DNA as described51.

#### Transcriptomic

##### Microarrays

50,000 CFSE-labelled OT-I cells were adoptively transferred in recipient mice subsequently infected with *Lm*-Ova N4, T4 or A8, and 3 days later 10,000 CFSE^low^ (divided) OT-I cells were flow-sorted based on congenic markers after enrichment for CD8^+^ T cells (using negative selection, as described above). Pelleted cells were stored in 700μl of TRIzol (Life Technologies) at -80C until RNA extraction. Total RNA was extracted using the RNAeasy Micro kit with RNase-Free DNase Set (Qiagen) according to the manufacturer protocol. The quality score and quantity of purified RNA was assessed with a Bioanalyzer RNA 6000 Pico Chip (Agilent). Total RNA was then converted to cDNA, amplified and hybridized to Affymetrix Mouse Transcriptome Array 1.0 Pico. Raw CEL files were preprocessed and normalized using Affymetrix Expression Console (version 1.4.1.46) and resulting data were analyzed with the Affymetrix Transcriptome Analysis Console (version 3.1.0.5). We calculated fold-differences between experimental groups and tested significance using one-way ANOVA (unpaired). Significantly up and down regulated genes were defined with at least a 1.5 fold expression difference and a *p*-value ≤ 0.05. Over-representation of biological process (BP) gene ontology (GO) terms was calculated using DAVID 6.8 and visualized in scatterplots using ggplot2 in R.

##### RNA-seq

1,000 memory OT-I cells were flow-sorted based on congenic markers after enrichment for CD8^+^ T cells (as described above). cDNA was synthesized and amplified directly from intact cells using SMART-Seq v4 Ultra Low Input RNA Kit for Sequencing (Takara Bio USA) according to the manufacturer protocol. The quality score and quantity of purified cDNA was assessed with Qubit dsDNA HS Assay kit (Life technologies) and Bioanalyzer High Sensitivity DNA assay (Agilent). Illumina cDNA library was prepared using Nextera XT DNA Library Preparation Kits according to manufacturer protocol. The quality score and quantity of purified cDNA library was assessed with Qubit dsDNA HS Assay kit (Life technologies) and Bioanalyzer High Sensitivity DNA assay (Agilent). The library samples were performed on Illumina HiSeq (2X150bp and dual index configuration) by Genewiz Inc. Reads were aligned to the Mouse reference mm10 using STAR aligner (v2.4.2a) ^52^. Quantification of genes annotated in Gencode vM5 were performed using featureCounts (v1.4.3) and quantification of transcripts using Kalisto. QC was collected with Picard (v1.83) and RSeQC ^53^(http://broadinstitute.github.io/picard/). Normalization of feature counts was done using the DESeq2 package, version 1.10.1. Differentially expressed genes were identify using negative binomial distribution as implemented in DESeq 2 (R package). Significantly up and down regulated genes were defined with at least a 1.5 fold expression difference and an adjusted *p*- value ≤ 0.05.

#### Epigenetic profiling

We performed ATAC-seq analysis on two biological replicates per group as previously described^54^. Briefly, 15,000-50,000 OT-I memory cells were flow-sorted from negatively-enriched splenocytes from host mice adoptively transferred with WT or *Il2ra^mut/mut^* OT-I cells and immunized with *Lm*-Ova N4 or T4. Then nuclei were isolated using a solution of 10 mM Tris-HCl, 10 mM NaCl, 3 mM MgCl2, and 0.1% IGEPAL CA-630. Immediately following nuclei isolation, the transposition reaction was conducted using Tn5 transposase and TD buffer (Illumina) for 45 min at 37° C. Transposed DNA fragments were purified using Qiagen Mini- Elute Kit and PCR amplified using NEB Next High Fidelity 2x PCR master mix (New England Labs) with dual indexes primers (Illumina Nextera). Genomic Alignment of sequencing reads; for all sequenced data, paired-end reads were trimmed for adaptors and removal of low-quality reads using Trim_galore (v0.3.7). Trimmed reads were mapped to the Mus Musculus genome (mm10 assembly) using Picard (v1.92). Reads were then filtered to exclude mitochondrial DNA or duplicates using samtools (v0.1.19). For peak calling, all positive-strand reads were shifted 4bp downstream to center the reads on the transposon binding event. We used MACS2 (v2.1.0) at a *p*-value of 0.01. Irreproducible discovery rate (IDR) calculations using scripts provided by the ENCODE project (https://www.encodeproject.org/software/idr/; v2.0.2 and v2.0.3) were performed on all pairs of replicates, keeping only reproducible peaks showing an IDR value of 0.05 or less. For peak annotation and analysis, peak assignment was done using ChipSeeker ^55^. Promoter regions were defined as peaks that overlapped a region that was +/- 3kb from the transcriptional start site (TSS). Intragenic (intronic and exonic) peaks were defined as any peak that overlapped with annotated intronic and exonic peaks, respectively, based on the annotation database. Intergenic peaks were defined as any non-promoter or non-intragenic peaks and were assigned to the gene of the nearest TSS based on the distance from the start of the peak. Conserved and unique open chromatin regions (OCRs) were defined using the bedTools intersect function (v2.29.1) at different level of comparison as represented in Figure 4B. First level of comparison allowed us to identify unique (non-overlapping) peaks between WT N4 and WT T4

(A); MUT N4 and MUT T4 (B); WT N4 and MUT N4 (X); WT T4 and MUT T4 (Y). Similar approach was used in a second level analysis where we identified conserved (overlapping) peaks between A and B (TCR controlled OCRs), overlapping peaks between X and Y (IL-2 controlled OCRs) and peaks unique of A, B, X, Y (OCRs controlled by both TCR and IL-2). The 3 master lists of peaks, TCR, IL-2 and TCR+IL-2, were used for further analysis after filtering for redundant peaks. De novo analysis and known-motif analysis on OCRs were performed using HOMER (v4.4) ^56^ with the findMotifsGenome.pl function. Likelihood of TF binding among unique peaks was evaluated using FIMO tool ^57^. Prior to motif analysis, for overlapping peaks (TCR and IL-2), peaks coordinates were adjusted to cover the full region represented by peak 1 and peak 2 using for each peak the extremes start and stop coordinates. To assess variability among the ATAC-seq datasets, we performed Pearson correlation across samples only considering overlapping peaks across conditions. Over-representation of BP GO terms was calculated using Panther and visualized in scatterplots using ggplot2 in R.

### Statistics

Statistical significance was calculated using an unpaired Student *t* test with GraphPad Prism software and two-tailed *p* values are given as: (*) *p*<0.1; (**) *p*<0.01; (***) *p*<0.001; (****) p<0.0001 and (ns) *p*>0.1. All *p* values of 0.05 or less were considered significant and are referred to as such in the text.

## Acknowledgments

We thank the Einstein FACS and genomic core facilities, and John Wherry (U Penn) for sharing his ATAC-seq protocol. **Funding**: This work was funded by the National Institute of Health Grants (NIH) AI103338, Hirschl Caulier Award to GL. LC received fellowships from ARC, Fondation Bettencourt-Schuller and the American Association of Immunology (AAI). SSC and EG were respectively supported by NIH training grant T32 AI070117 and NIH MSTP training grant T32 GM7288. Core resources for FACS were supported by the Einstein Cancer Center (NCI cancer center support grant 2P30CA013330).

## Author contribution

SSC designed, performed and interpreted a majority of experiments, assembled many Figures, and contributed to writing/editing of the manuscript. EG conducted and analyzed multiple functional experiments including all revisions, replotted/analyzed data and contributed to figures and editing of the manuscript. LC laid the ground work for the project, performed and interpreted many experiments. SA and GAB conducted and analyzed the *in vitro* quantitative analyses of OT-I cell priming. FA, SO and DZ provided key recombinant *Lm* strains. FD with GL, SSC and EG, conducted, guided and interpreted analysis of all transcriptomic and epigenetic data. GL designed and interpreted experiments with LC, SSC, EG and all other authors, contributed to Figure design and editing, and wrote the paper. **Competing interests**: The authors declare that no competing interests exist. **Data materials and availability:** The accession number for the ATAC-seq and RNA-seq data reported in this paper is GEO: GSE152394. All data is available in the main text or the supplementary materials.

## Supplementary Materials

**Figure S1.**
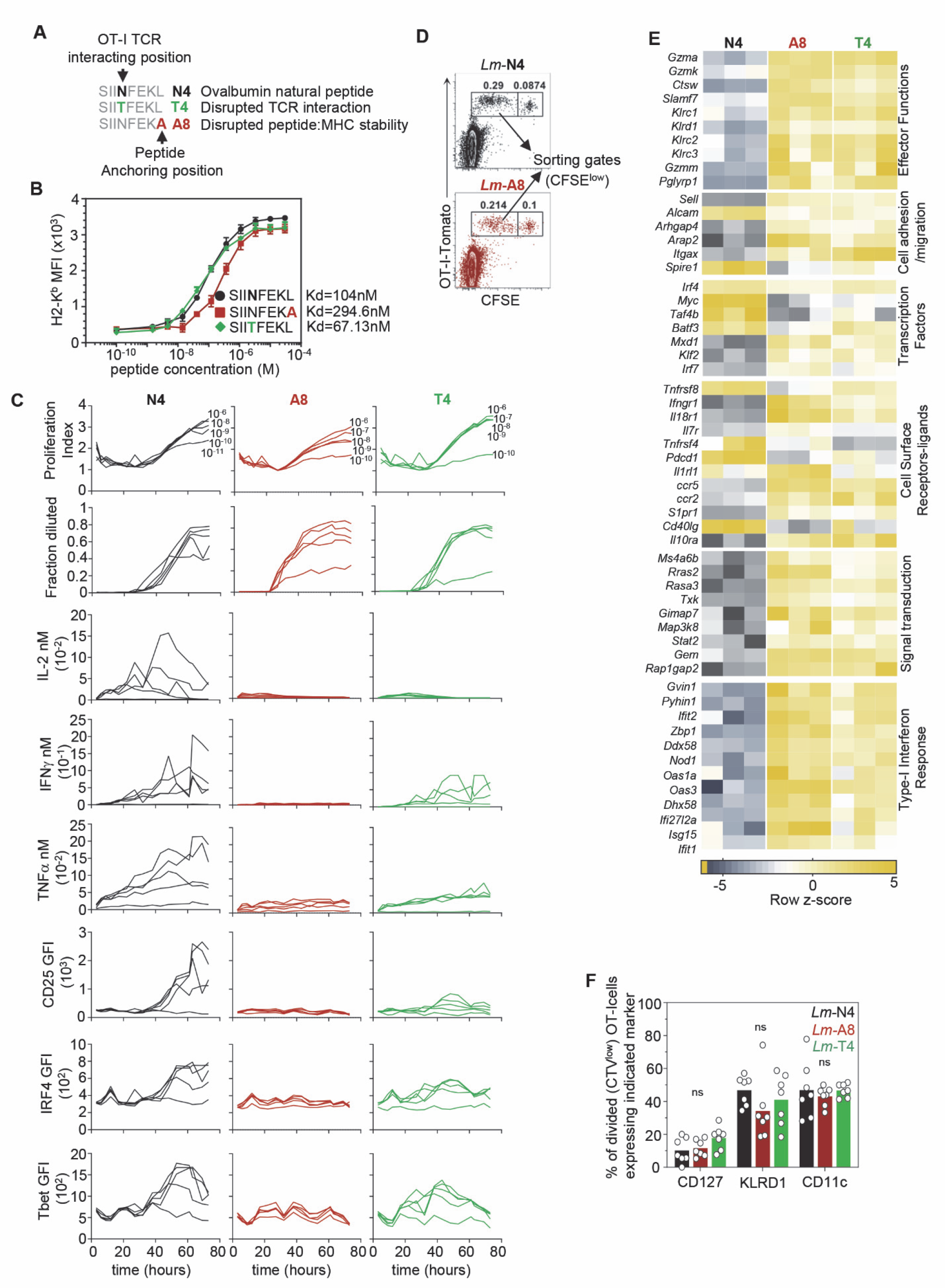
TCR signaling strength affects early CD8^+^ T cell activation program. (**A**) Ova-derived SIINFEKL and its APLs. (**B**) H2-K^b^ RMA-S stabilization assay comparing Ova N4, A8 and T4. Calculated Kd for each peptide is shown. (**C**) Graphs show *in vitro* stimulation of OT-I cells cultured with various concentrations of Ova N4, A8 and T4. (**D**) Representative FACS dot plots showing CFSE^low^ N4- and A8-primed OT-I cells 3 days post-infection subsequently sorted for microarray analysis. (**E**) Heat map of selected genes grouped under the indicated categories and for which expression is significantly different between N4-, A8- and T4-primed OT-I cells. (**F**) Proportion of CTV^low^ (divided) OT-I cells expressing indicated markers 3 days post immunization with *Lm-*Ova N4, A8 or T4.

**Figure S2.**
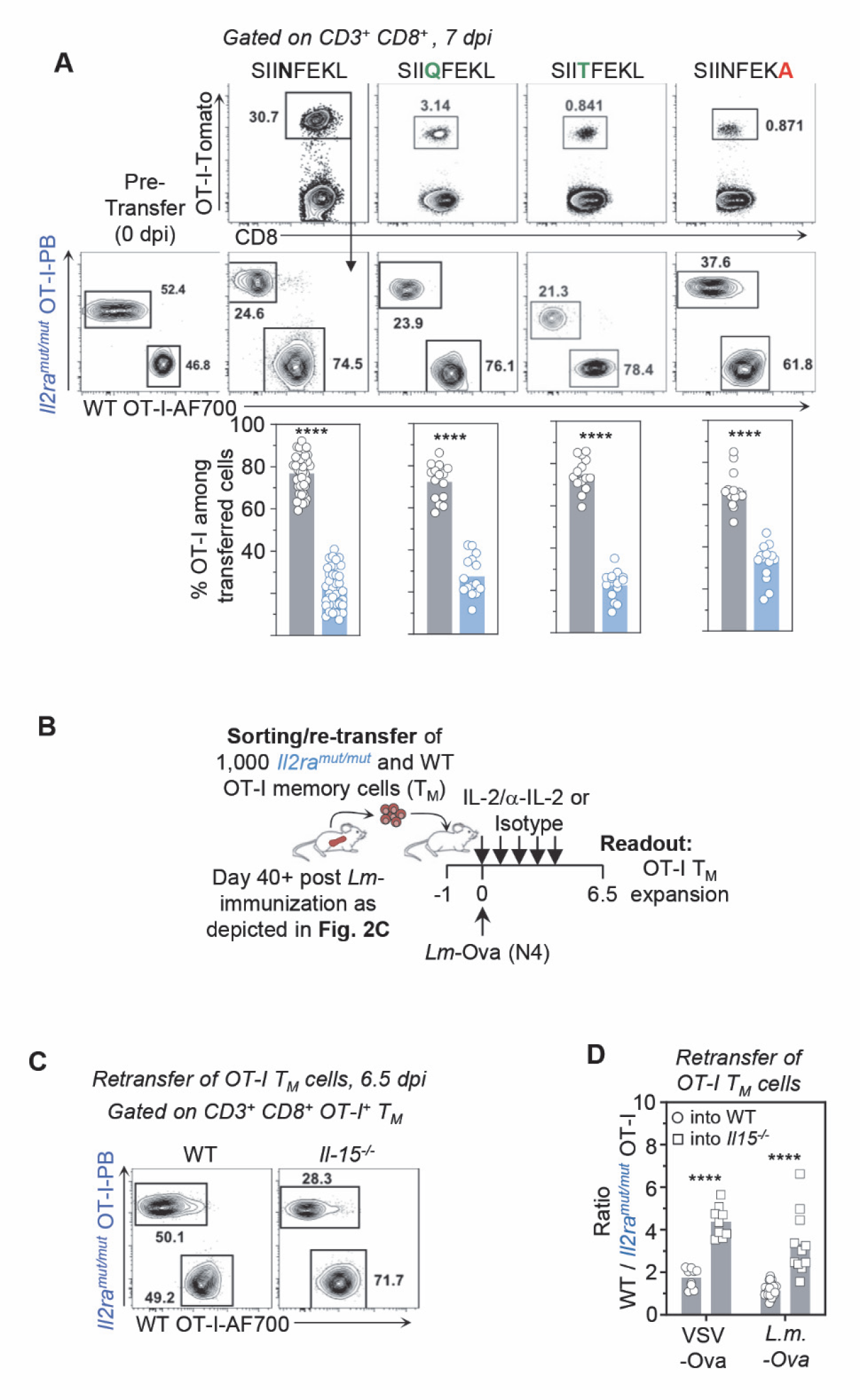
CD8^+^ T cells primed with weak TCR and IL-2 signals failed to expand competitively during primary challenge. (**A**) Representative FACS dot plots of adoptively transferred *Il2ra^mut/mut^* versus WT OT-I cells 7 days post primary infection with *Lm* expressing N4, Q4, T4 or A8. Bar graphs show the relative pooled frequency of expanded OT-I cells across >5 replicate experiments (n=14-42 mice). (**B**) Schematic of experimental design complementing Figure 2C. (**C**) Representative FACS dot plots of relative proportions of *Il2ra^mut/mut^* and WT OT-I memory cells from Figure 2E. (**D**) Bar graph shows the ratio of WT versus *Il2ra^mut/mut^* OT-I memory cells adoptively transferred in either WT or *Il15^-/-^* recipient mice subsequently infected with indicated pathogens. In all panels, each symbol represent 1 mouse and *p*-values are indicated when relevant with \**p* < 0.05; \*\**p*< 0.01; \*\*\**p*< 0.001; \*\*\*\**p*< 0.0001; NS, not significant, using two-tailed unpaired Student’s *t* test.

**Figure S3.**
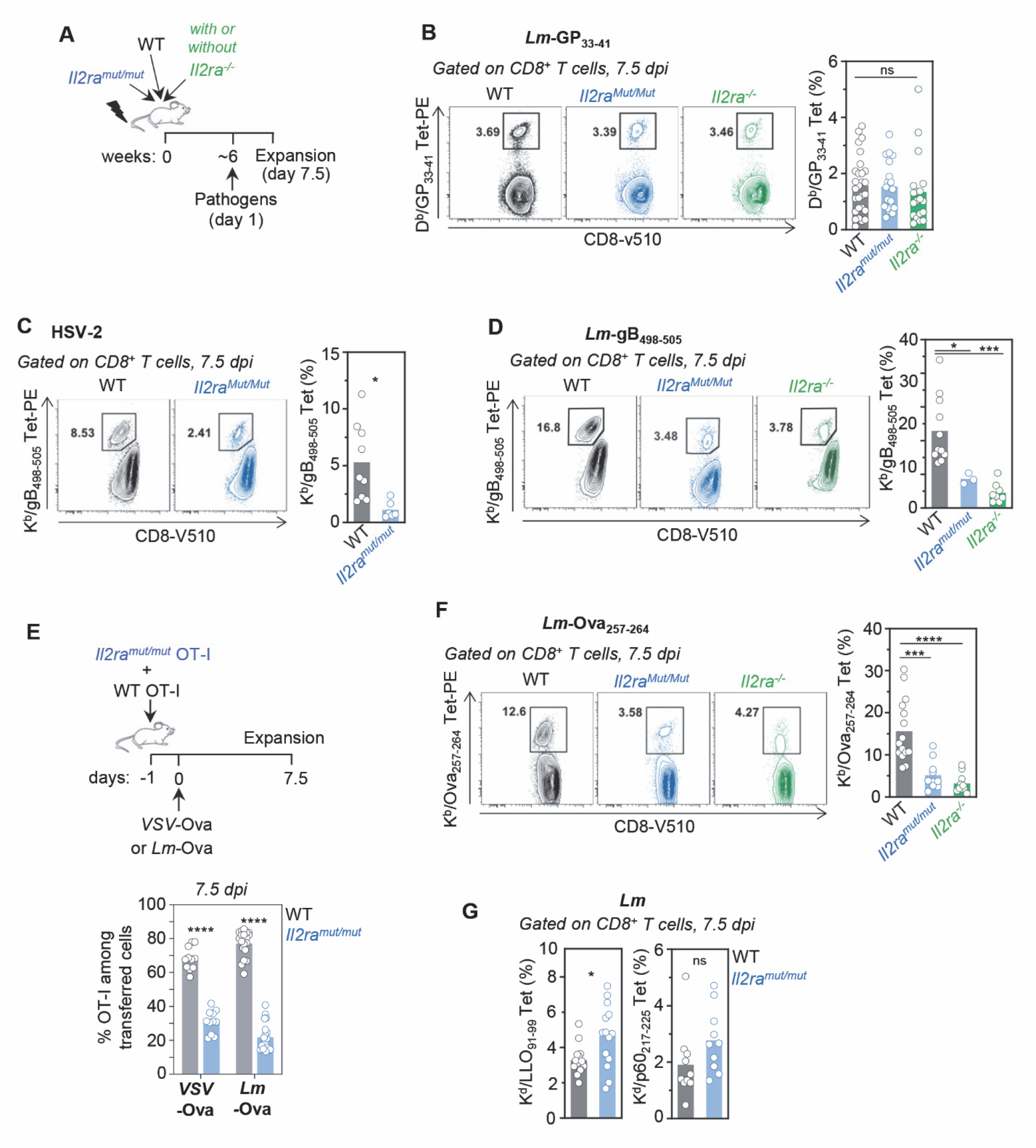
Analysis of WT and IL-2-disrupted pathogen-specific CD8^+^ T cell primary responses in response to various epitopes and challenge infections. **A** Schematic of BM chimera experimental setup. Irradiated hosts were reconstituted with BM harvested from WT, *Il2ra^mut/mut^* and/or *Il2ra^-/-^* mice (ratio 1:1:1), and 6 weeks later, infected with indicated pathogens. **(B)** Representative FACS plots of WT and *Il2ra^mut/mut^* GP33-41/D^b^ Tet^+^ CD8^+^ T cells in BM chimeras infected by *Lm*-GP33-41. Bar graphs show the relative frequencies of *Il2ra^mut/mut^* versus WT D^b^/GP33-41-specific Tet^+^ CD8^+^ T cells 7.5 days post primary infection in 2-4 replicate experiments (n=4-20 mice). **(C)** Representative FACS dot plots of WT and *Il2ra^mut/mut^* gB498- 505/K^b^ Tet^+^ CD8^+^ T cells in BM chimeras infected by HSV-2. Bar graph shows the relative frequency of *Il2ra^mut/mut^* versus WT Tet^+^ CD8^+^ T cells 7.5 days post primary infection across 2 replicate experiments (n=7-10 mice). **(D)** Representative FACS dot plots of WT and *Il2ra^mut/mut^* gB498-505/K^b^ Tet^+^ CD8^+^ T cells from BM chimeras infected by *Lm*-gB498-505. Bar graph shows the relative frequency of *Il2ra^mut/mut^* versus WT Tet^+^ CD8^+^ T cells 7.5 days post primary infection across 2 replicate experiments (n=3-11 mice). **(E)** Schematic of experimental setup to examine primary expansion of adoptively transferred *Il2ra^mut/mut^* and WT OT-I cells. Bar graph shows the relative frequency of *Il2ra^mut/mut^* and WT OT-I cells 7.5 days post primary infection across 2-3 replicate experiments (n=3-11 mice). **(F)** Representative FACS dot plots of Ova257-264/K^b^ Tet^+^ WT and *Il2ra^mut/mut^* CD8^+^ T cells in BM chimeras infected by *Lm*-Ova257-264. Bar graph shows the relative frequency of *Il2ra^mut/mut^* versus WT Tet^+^ CD8^+^ T cells 7.5 days post primary infection across 2-3 replicate experiments (n=10-15 mice). **(G)** Bar graphs show the relative frequency of *Il2ra^mut/mut^* versus WT indicated Tet^+^ CD8^+^ T cells 7.5 days post primary *Lm* infection of B6-K^d^ mice across 2 replicate experiments (n=10-15 mice).

**Figure S4.**
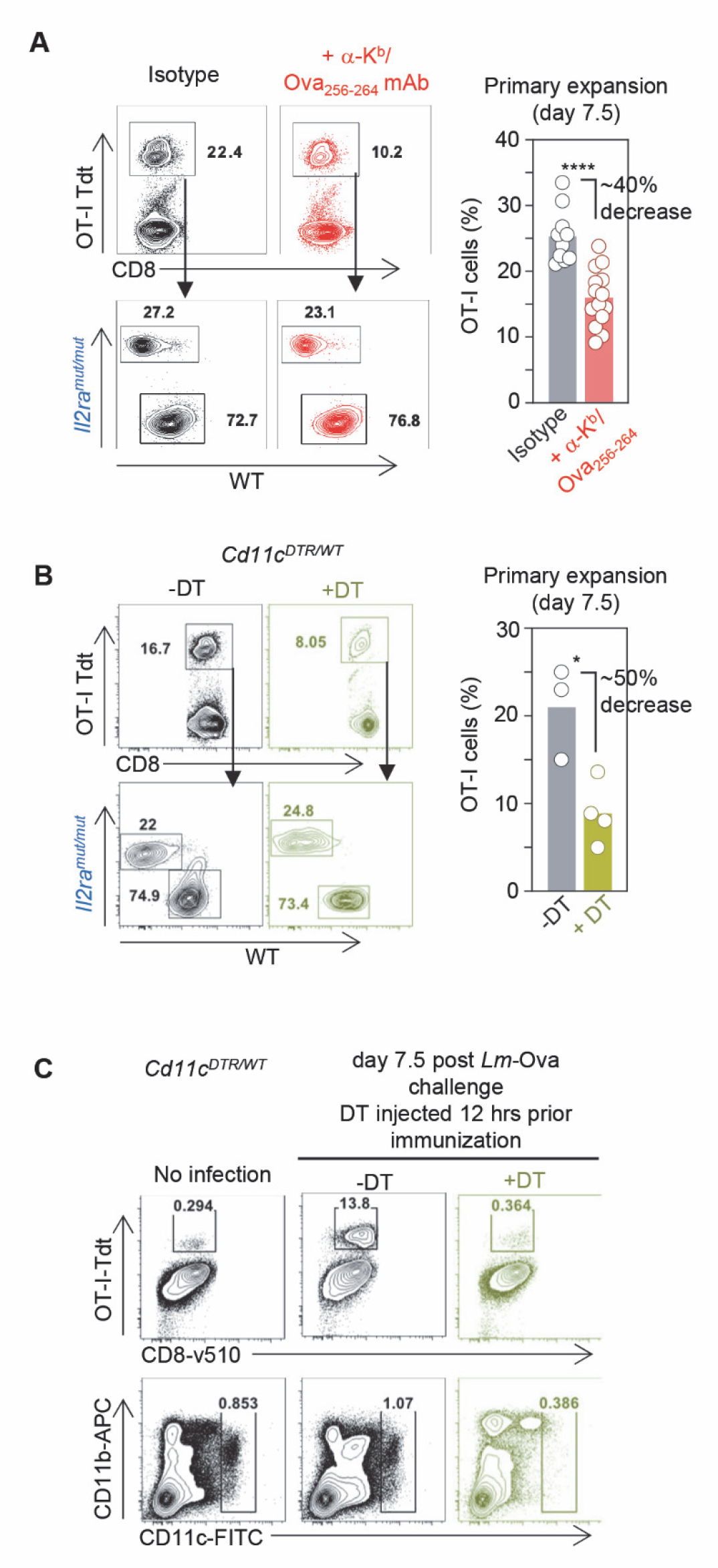
Disrupting antigen presentation impairs primary CD8^+^ T cell responses. *Il2ra^mut/mut^* and WT OT-I cells adoptively transferred to naïve WT (**A**) or *Cd11c^DTR/WT^* (**B, C**) mice were subsequently infected the day after with *Lm*-OvaN4. Antigen presentation was disrupted or not (isotype) by injecting (**A**) anti-MHC-I K^b^/Ova256-264 mAb or (**B**) diphtheria toxin (DT) 48 hrs post-infection and spleen cells were stained for FACS analysis of OT-I cell expansion of each genotype for the various treatments 7.5 days later. Representative FACS dot plots are shown. Bar graph shows the frequency OT-I cell expansion in anti-K^b^/Ova256-264 mAb versus isotype-treated or DT-treated versus untreated mice in 2-3 replicate experiments (n=3-10 mice). (**C**) Same design as in (B), but with *Il2ra^mut/mut^* and WT OT-I cells adoptively transferred to *Cd11c^DTR/WT^* recipient mice further injected with diphtheria toxin (DT) 12 hours prior infection to deplete CD11c^+^ cells. Dot plots show OT-I cell expansion (or lack of) and CD11c^+^ DC depletion 7.5 days post challenge infection in 1 of 4 representative mice.

**Figure S5.**
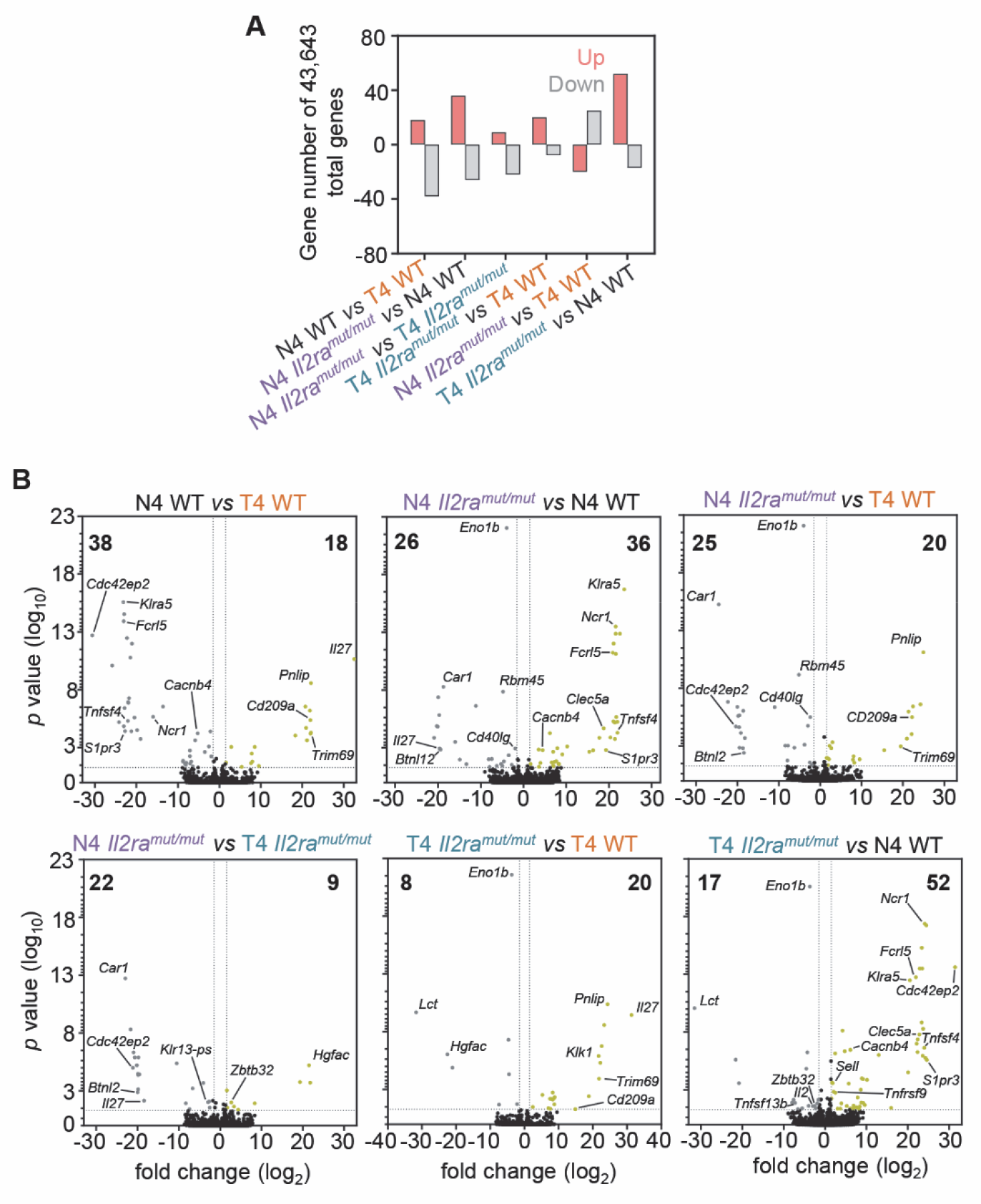
The strength of TCR and/or IL-2 signaling only minimally affects gene expression in resting memory CD8^+^ T cells. **A** Bar graph shows the number of significantly up- and down-regulated genes in *Il2ra^mut/mut^* and WT resting OT-I memory cells primed after infection *Lm*-OvaN4 or *Lm*-OvaT4, defined as genes with at least 1.5 fold change, adjusted *p* value ≤ 0.05 in each respective comparison. **(B)** Volcano plot of *p*-values versus gene expression fold change in each respective comparisons. Significantly up- and down-regulated genes were defined as genes with at least 1.5 fold change, p-value ≤ 0.05, and colored gold or gray, respectively.

**Figure S6.**
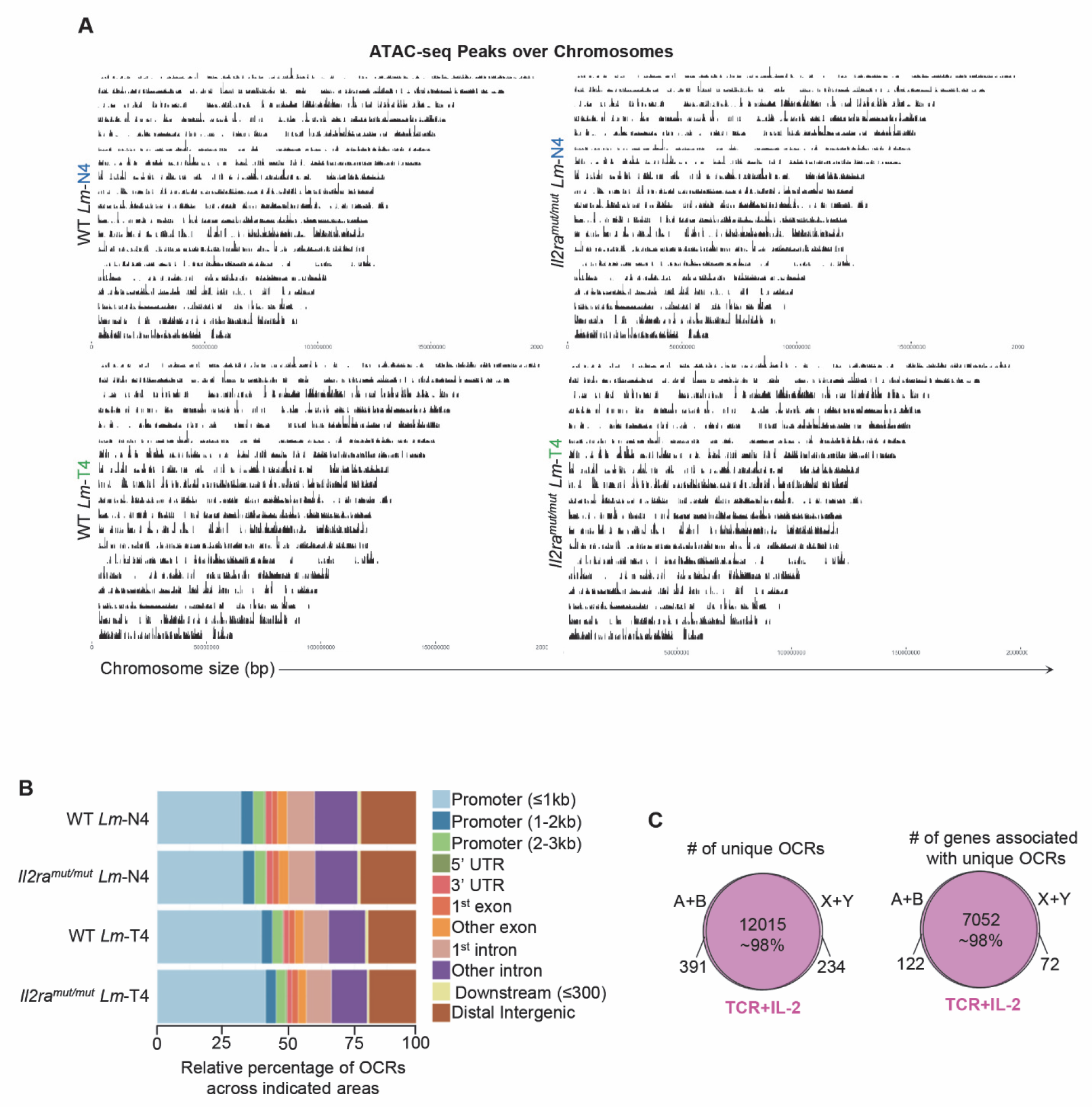
Analysis of chromatin remodeling in OT-I memory cells primed under the various conditions. **A** Distribution of ATAC-seq peaks across chromosomes in the various resting OT-I memory cells compared. **(B)** Relative OCR proportions across genome areas in *Il2ra^mut/mut^* and WT resting OT-I memory cells primed after infection *Lm*-OvaN4 or *Lm*-OvaT4. **(C)** Venn diagrams comparing the number of unique OCRs and the number of genes associated with these unique OCRs induced by modulation of TCR+IL-2 signals.

**Figure S7.**
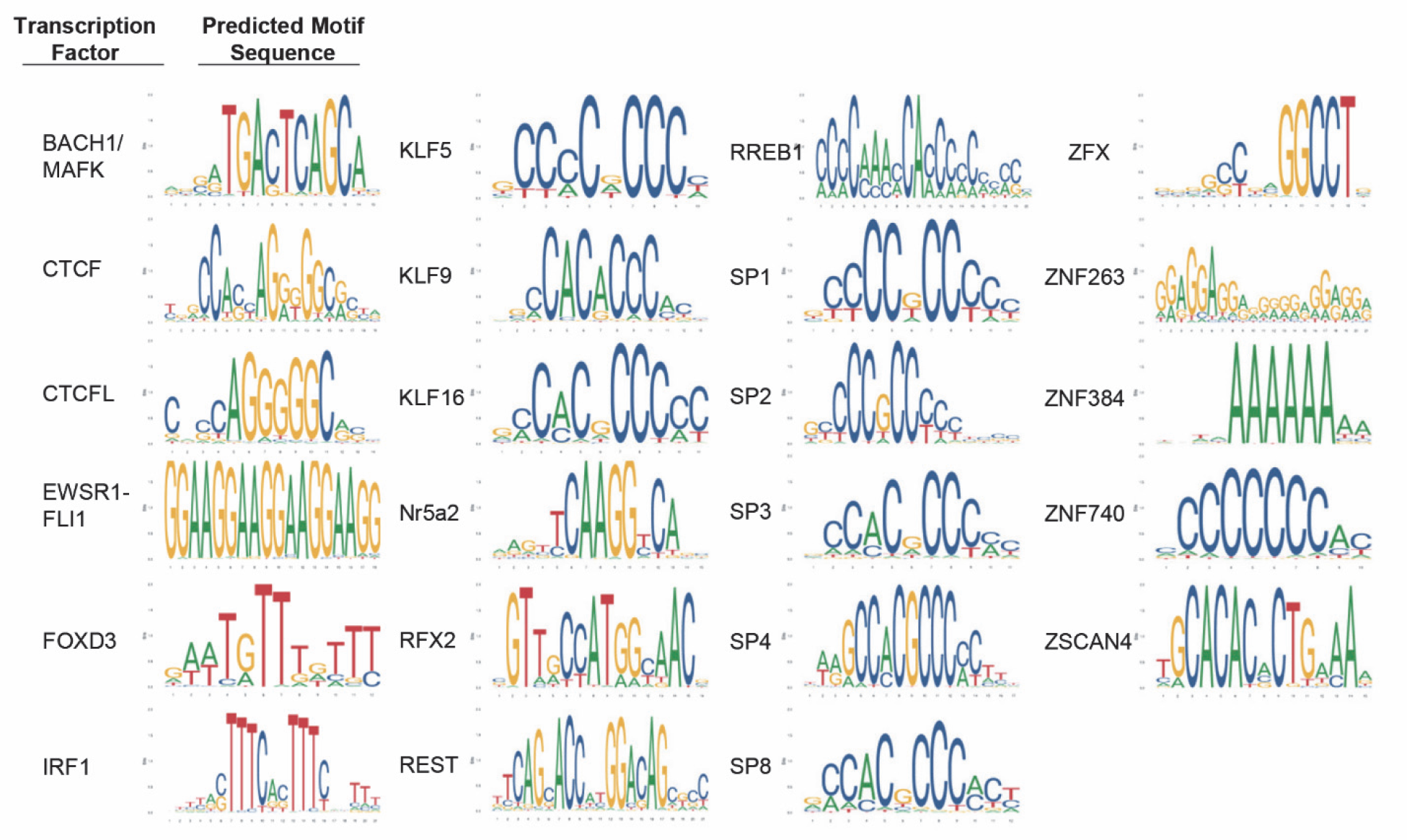
DNA binding motifs for indicated transcription factors. The JASPAR database was used to recover the predicted DNA binding sequences for listed TFs revealed from the FIMO analysis of the differentially accessible OCRs of Figure 7A genes.

**Figure S8.**
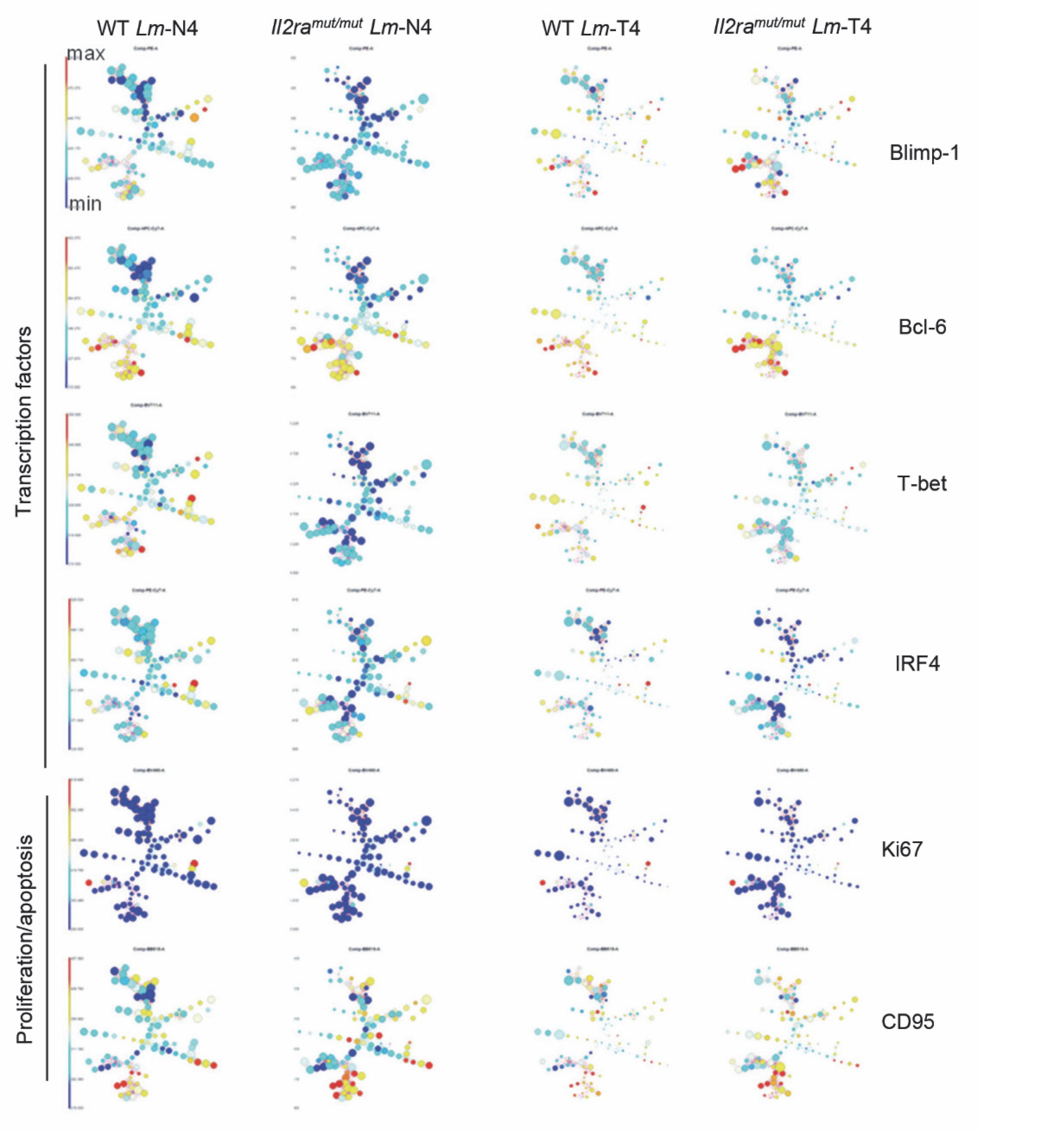
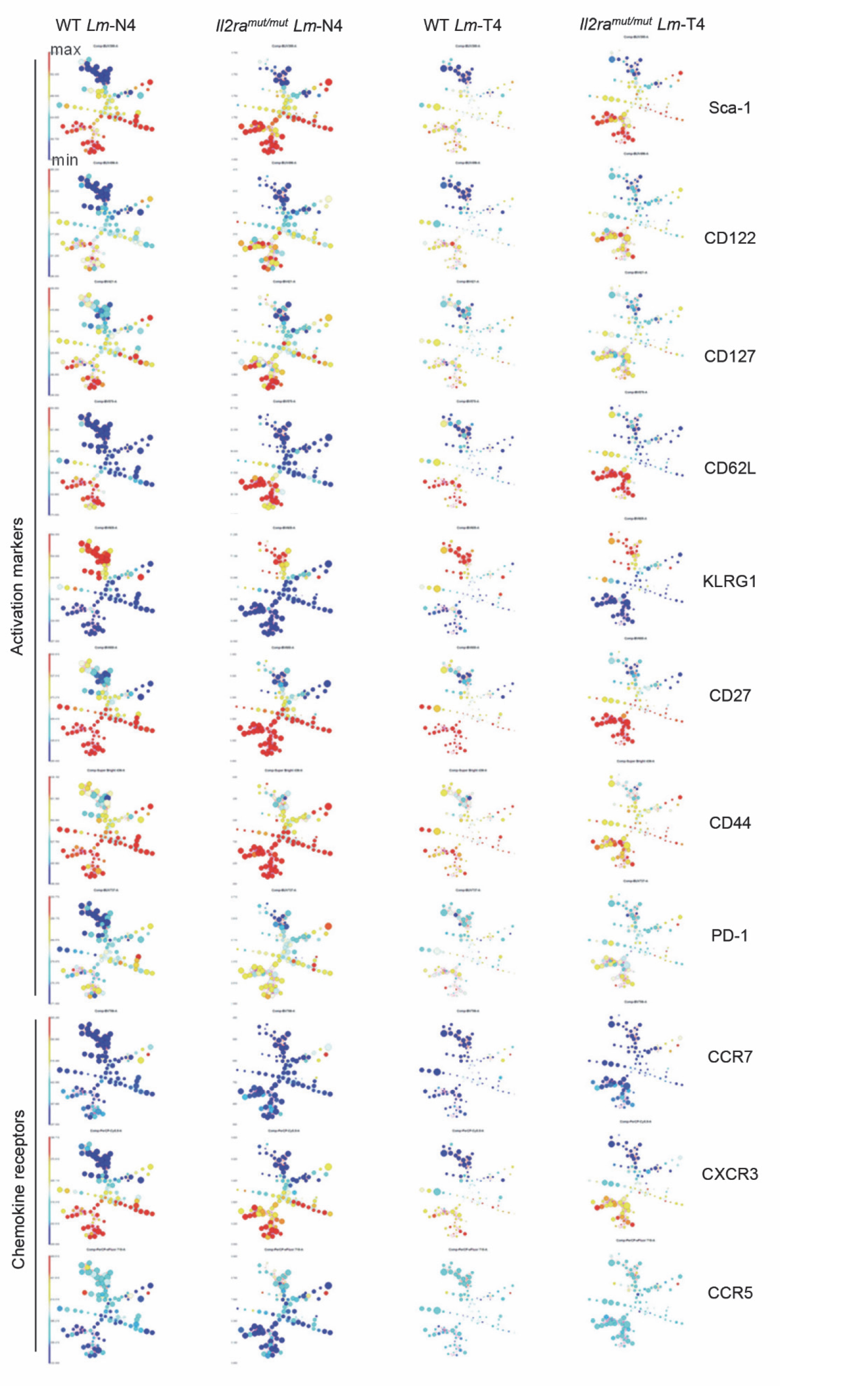
FlowSOM analysis of *Il2ra^mut/mut^* and WT OT-I memory cells primed with either *Lm*-N4 or *Lm*-T4 after staining with a panel of 26 memory cell-relevant markers. The expression level of each marker within each node is represented in a color scale.

**Figure S9.**
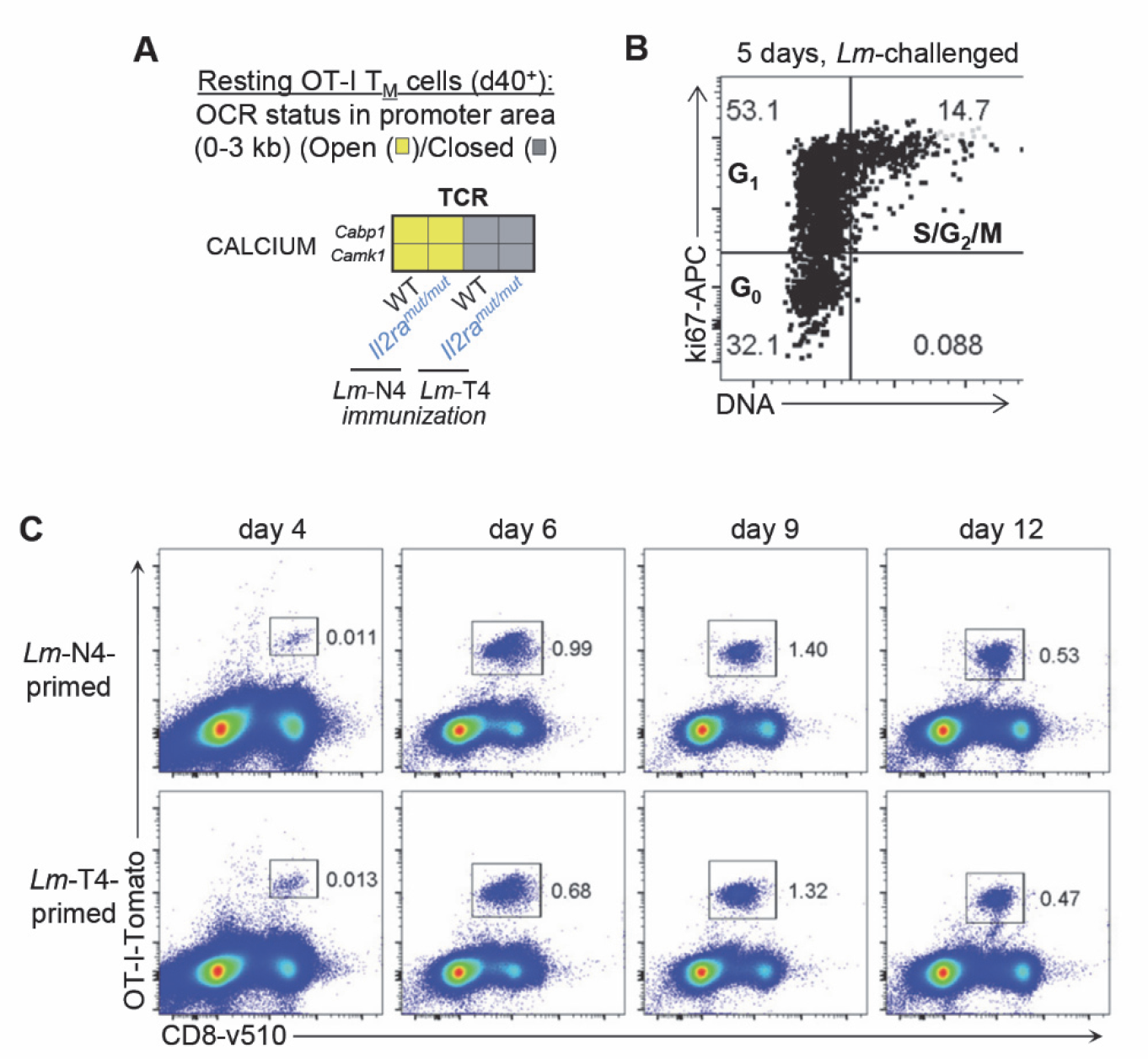
Functional analysis of reactivated memory CD8^+^ T cells. (**A**) Heat map shows OCRs (yellow) in genes encoding calcium fluxes that were revealed in Figure 6B when only TCR signals were modulated, in the promoter area and close to the TSS (+/- ∼3kb). Chromatin region status (open, yellow/closed, grey) in WT and *Il2ra^mut/mut^* OT-I memory cells primed with *Lm*-N4 or *Lm*-T4 are reported. (**B**) Representative FACS dot plot of cell cycle analysis of splenic CD8^+^ T cells 5 days post *Lm*-immunization, after staining for cell surface CD3, CD8 and intracellular Ki67 and DNA. (**C**) Experimental design is that of Figure 9A, “late reactivation”. 40 days post primary infection, 1,000 *Il2ra^mut/mut^* and WT OT-I memory cells from each priming condition, were FACS-sorted and transferred at a 1:1 ratio to new hosts subsequently infected with *Lm*-OvaN4 the day after. At indicated days, spleen cells were stained to quantify frequencies of global OT-I memory cell expansion among CD8^+^ T cells. Data is a representative dot plot of n=15 mice.

**Figure S10.**
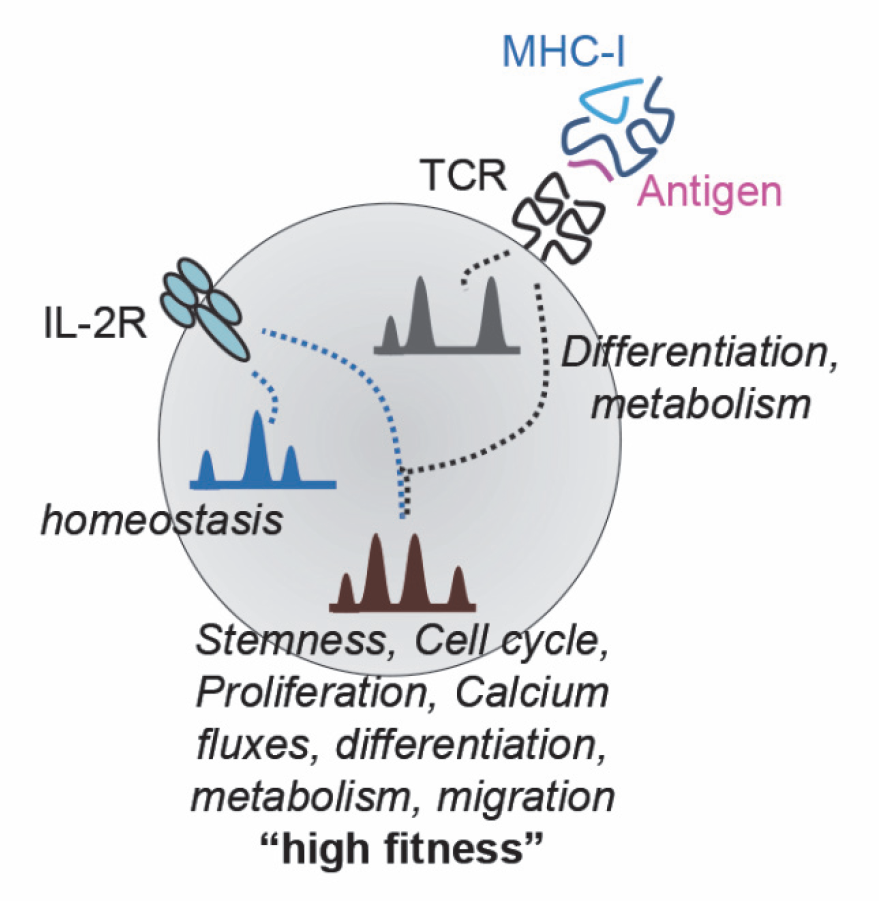
Proposed working model.

Table S1. GO pathways for Fig. 1C

Table S2. Transcriptomic analysis of resting OT-I memory cells for Figure 5A-S5

Table S3. OCR TCR, IL-2 and TCR+IL-2Table S3: OCRs

Table S4. GO pathways ATAC-seq comparisons for Figure 6D

Table S5: OCRs in promoters of selected genes and FIMO TF binding analysis for Figure 7

Table S6. Cytek FACS panel

Table S7. Table for antibodies used

## Notes

### Competing Interest Statement

The authors have declared no competing interest.

